# Simulating single-cell metabolism using a stochastic flux-balance analysis algorithm

**DOI:** 10.1101/2020.05.22.110577

**Authors:** David S. Tourigny, Arthur P. Goldberg, Jonathan R. Karr

## Abstract

Stochasticity from gene expression in single cells is known to drive metabolic heterogeneity at the level of cellular populations, which is understood to have important consequences for issues such as microbial drug tolerance and treatment of human diseases like cancer. Despite considerable advancements in profiling the genomes, transcriptomes, and proteomes of single cells, it remains difficult to experimentally characterise their metabolism at genome-scale. Computational methods could bridge this gap toward a systems understanding of single-cell biology. To address this challenge, we developed stochastic simulation algorithm with flux-balance analysis embedded (SSA-FBA), a computational framework for simulating the stochastic dynamics of the metabolism of individual cells using genome-scale metabolic models with experimental estimates of gene expression and enzymatic reaction rate parameters. SSA-FBA extends the constraint-based modelling formalism of metabolic network modelling to the single-cell regime, enabling simulation when experimentation is intractable. We also developed an efficient implementation of SSA-FBA that leverages the topology of embedded FBA models to significantly reduce the computational cost of simulation. As a preliminary case study, we built a reduced single-cell model of *Mycoplasma pneumoniae*, and used SSA-FBA to illustrate the role of stochasticity on the dynamics of metabolism at the single-cell level.

**SIGNIFICANCE:** Due to fundamental challenges limiting the experimental characterisation of metabolism within individual cells, computational methods are needed to help infer the metabolic behaviour of single cells from information about their transcriptomes and proteomes. In this paper, we present SSA-FBA, the first systematic framework for modelling the stochastic dynamics of single cells at the level of genome-scale metabolic reaction networks. We provide a robust and efficient algorithm for simulating SSA-FBA models, and apply it to a case study involving the metabolism, RNA and protein synthesis and turnover of a single *Mycoplasma pneumoniae* cell.

## INTRODUCTION

Describing the phenotypic behaviour of single cells is critical for a deeper understanding of biological tissues, organisms and populations. Recent experimental advances are driving a data explosion in systems biology by enabling researchers to profile multiple dimensions of single cells, including their genome, transcriptome, and proteome (1–3). Such single-cell measurements can yield information about thousands of individual cells in a single experiment. This can provide insights on intracellular function and the role of intercellular heterogeneity in a variety of biological systems ranging from microbial populations (4, 5) to human diseases such as cancer (6, 7).

Although metabolism is a key aspect of cellular physiology, methodologies for probing the metabolism of single cells are comparatively immature (8–10, 12, 13). This presents a barrier to studying a variety of phenomena such as metabolic reprogramming in tumours, which is now understood to be a central hallmark of cancer (14, 15). It is challenging to experimentally probe the metabolism of single cells due to low abundances of many metabolites, the compartmentalisation of eukaryotic cells, and the wide structural diversity of metabolites (10). Moreover, faster timescales of enzymatic reactions makes measuring the dynamics of metabolism more difficult than measuring that of DNA replication or gene expression. These experimental obstacles to studying single-cell metabolism necessitate the development of computational techniques that can infer the metabolism of single cells from other information, such as single-cell transcriptomics or proteomics and population-level metabolic data (11).

While our current capabilities to probe and model the metabolism of single-cells are limited, considerable attention has been devoted to the metabolism of cellular populations. For example, metabolic network modelling has achieved a great deal of success combining limited experimental data and computational simulation to study gene essentiality and guide metabolic engineering (16–18). Extensions of these population-based metabolic network modelling frameworks, such as dynamic metabolism expression models (19) and dynamic enzyme-cost flux-balance analysis (20), have also been developed to incorporate the dynamics of gene expression. However, these deterministic approaches cannot capture effects that are relevant to the metabolism of single cells, where stochasticity is understood to play a pivotal role (21, 22). To date, there have only been a handful of attempts (see (23) and references therein) to extend metabolic network modelling to the single-cell regime. These efforts have mainly focused on integrating single-cell transcriptomics data with flux-balance analysis (FBA) in the context of cancer. Critically, these studies have not addressed the temporal dynamics of single cells, which is a key feature of their behaviour.

The existing approaches to stochastic modelling of single-cell metabolism can be organised into two categories: (a) (semi-)analytical treatment of rigorous models of individual metabolic pathways involving the expression of one (24, 25) or several (26, 27) enzymes catalysing a handful of metabolic reactions, or (b) empirical simulation of whole-cell models (28, 29) involving hybrid methods that combine ordinary differential equations (ODEs), particle-based stochastic simulation algorithms (SSA, (30, 31)), and dynamic FBA (DFBA, (32)). Although the shared aim of these approaches is to relate single-cell behaviour to genotype, the two categories fall at opposite ends of a wide spectrum: whole-cell modelling aims to accommodate as much detail as possible, but no framework is yet available to rigorously simulate entire cells. On the other hand, analytical methods that can provide a mechanistic understanding of small pathways are intractable for entire cells. Furthermore, existing exact, approximate and hybrid stochastic simulation methods for large-scale biochemical networks (reviewed in (33)) remain a long way from applicability to single-cell metabolism because, unlike FBA, they still rely on explicitly encoding the dependence of reaction rate functions on enzymatic kinetic parameters. Here, we introduce a stochastic extension of FBA that we call stochastic simulation algorithm with flux-balance analysis embedded (SSA-FBA), which enables systematic systems-scale simulations of the metabolism of single cells. We believe SSA-FBA is a powerful computational tool for simulating stochastic dynamics of metabolic network models at the single-cell level.

The remainder of this paper is organised as follows. We first outline the concepts of SSA-FBA and relate it to a formal description of single-cell metabolism based on the chemical master equation. Next, we compare exact and approximate implementations of SSA-FBA, and then introduce an advanced algorithm that significantly improves the efficiency of exact simulations. As a case study, we subsequently present simulation results of an SSA-FBA model of a reduced single *Mycoplasma pneumoniae* cell representing 505 reactions where, as an illustrative example, we can explore the consequences of stochasticity for the dynamics of adenosine triphosphate (ATP) production and consumption at the single-cell level. We conclude the paper with a discussion of the main results and directions for future work.

## METHODS

We implemented an SSA-FBA simulation package in C++ and Python. Further details on SSA-FBA, its implementation and the case study can be found in the Supplementary Appendix, and all code and data are freely available open-source at https://gitlab.com/davidtourigny/single-cell-fba.

## RESULTS

### Stochastic simulation algorithm with flux-balance analysis embedded

Stochasticity in the metabolism and growth of single cells is generally believed to emerge primarily from fluctuations in enzyme expression levels (21, 22, 24, 25, 34) (see Figure 1, which provides an overview of sources and consequences of stochasticity at the single-cell level). Due to the relatively high copy numbers of most metabolites, metabolism is thought to have little intrinsic stochastic variation compared to gene expression (21). This observation motivates SSA-FBA as an appropriate framework for modelling single-cell metabolism, where the dynamics of reaction fluxes internal to a metabolic network are captured deterministically by FBA, while SSA is used to model changes in the copy numbers of enzyme molecules and metabolites that are produced or consumed on the periphery of the metabolic network.

**Figure 1:**
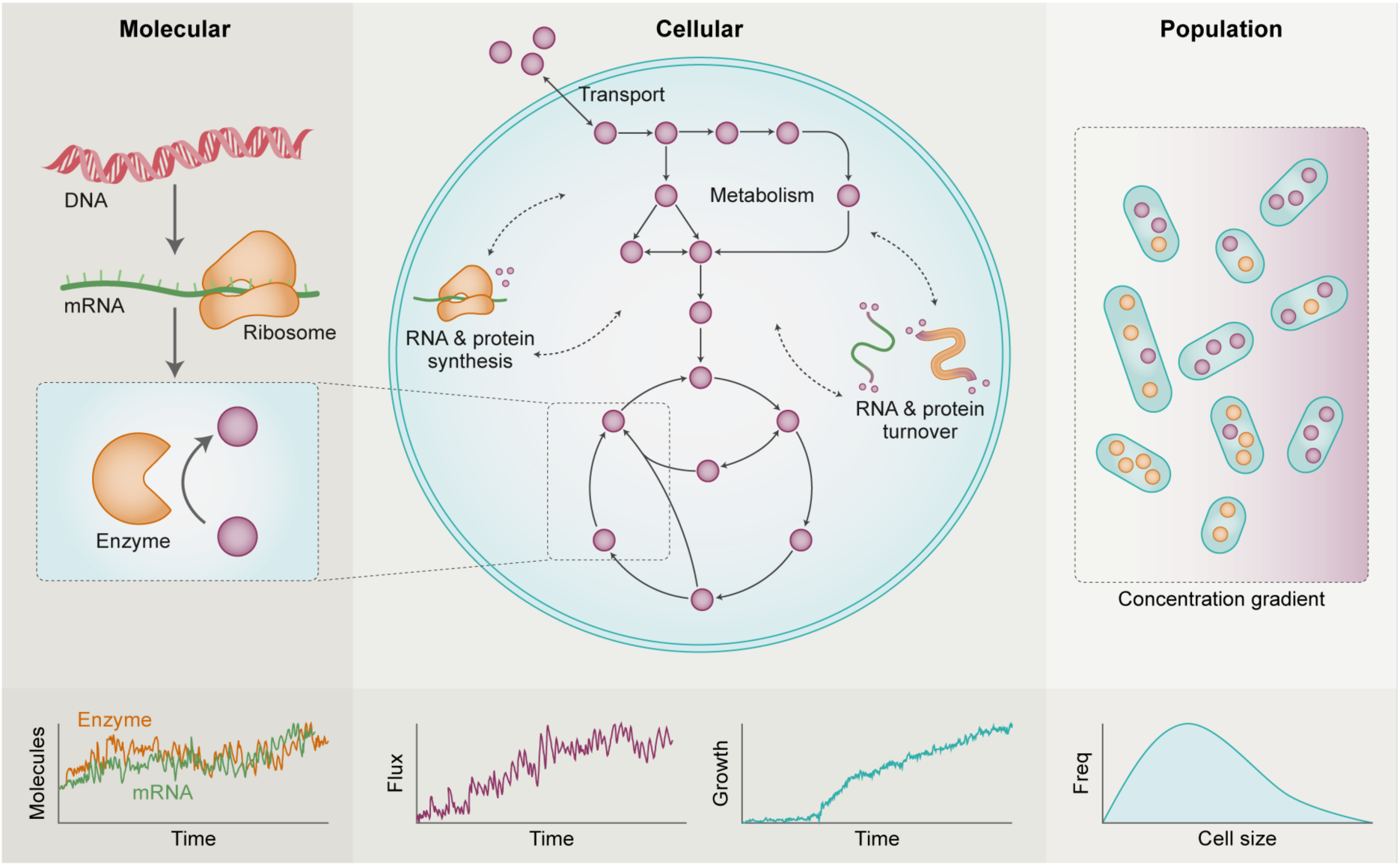
The metabolic behaviour of single cells is influenced by the spatiotemporal dynamics of metabolites and enzymes at multiple scales. Temporal variation in the metabolic behaviour of single cells is primarily driven by fluctuations in gene expression and amplified or attenuated by their metabolic networks. For example, the structure of the reaction network combined with fluctuating enzyme and metabolite levels determine intracellular metabolic fluxes that, in turn, govern the dynamics of energy supply and molecular compositions of individual cells. The latter determines a total cell mass, which, assuming typical cell density, can be used to infer cell volume or size. The resulting stochastic nature of these properties at the single-cell level drives phenotypic heterogeneity modelled as a distribution over random variables at the population level, which can also be influenced by external factors such as environmental conditions.

The combined single-cell network of *M* metabolic and enzyme expression reactions can be captured by the chemical master equation (CME) (35)

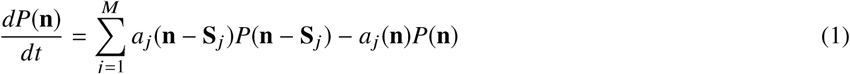

where **n** is an *N*-dimensional vector for counts of *N* chemical species, *a*_*j*_(**n**) is the propensity value of reaction *j* given **n**, and **S**_*j*_ is the stoichiometry of reaction *j*. The probability density function *P*(**n**) describes the probability of the system to occupy state **n**(*t*) at time *t*. Because CMEs are usually too complex to be solved directly, investigators often use SSA to sample trajectories through the state space of the CME (30, 31). One limitation of SSA is its computational cost, which is particularly acute for large chemical reaction networks. Various performance enhancements have therefore been developed to improve the run time of SSA. Since the average counts of enzymes are many times lower than those of metabolites, and metabolic reactions operate on a much faster time-scale than that of gene synthesis and degradation, several researchers have used *stochastic quasi-steady state assumptions* (stochastic generalisations of the quasi-steady state assumption in DFBA (32)) to approximate SSA trajectories at a lower computational cost. These methods correspond to a reduction of the CME (1) on the basis of time-scale separation or molecular abundances (see (36–39) and Supplementary Appendix S1).

While the above time-scale or abundance separation methods could in principle be used to simulate single-cell metabolism, investigators rarely have sufficient data to construct detailed dynamical models of metabolism in single cells due to the experimental challenges outlined in the Introduction. Instead, we realised that the constraint-based formalism of (D)FBA (16, 17, 32) could be used to identify a numerical solution to the deterministic quasi-steady state conditions of the metabolic reaction network with a small amount of data at a modest computational cost. In short, SSA-FBA embeds an FBA model into SSA, analogous to the way that an FBA model is embedded into a system of ODEs in DFBA. Marginalisation of the CME over copy numbers of metabolites internal to the metabolic reaction network motivates describing the dynamics of macromolecules and metabolites external to the metabolic reaction network using SSA, but where propensity values for metabolic reactions are obtained by solving the embedded FBA problem. In turn, the embedded FBA problem and therefore the resulting propensity values depend on the counts of macromolecules and external metabolites, which implies the embedded FBA problem must be dynamically updated and resolved for metabolic propensity values over the course of simulation. This embedding enables SSA-FBA to simulate the stochastic behaviour of the metabolism of single cells analogous to how DFBA enables deterministic simulation of the dynamic behaviour of the metabolism of populations.

First, SSA-FBA separates reactions into three mutually disjoint subsets based on whether the species that participate in the reaction are defined to be internal or external to the metabolic reaction network (Figure 2A). The three subsets of reactions in an SSA-FBA model are as follows:

**Figure 2:**
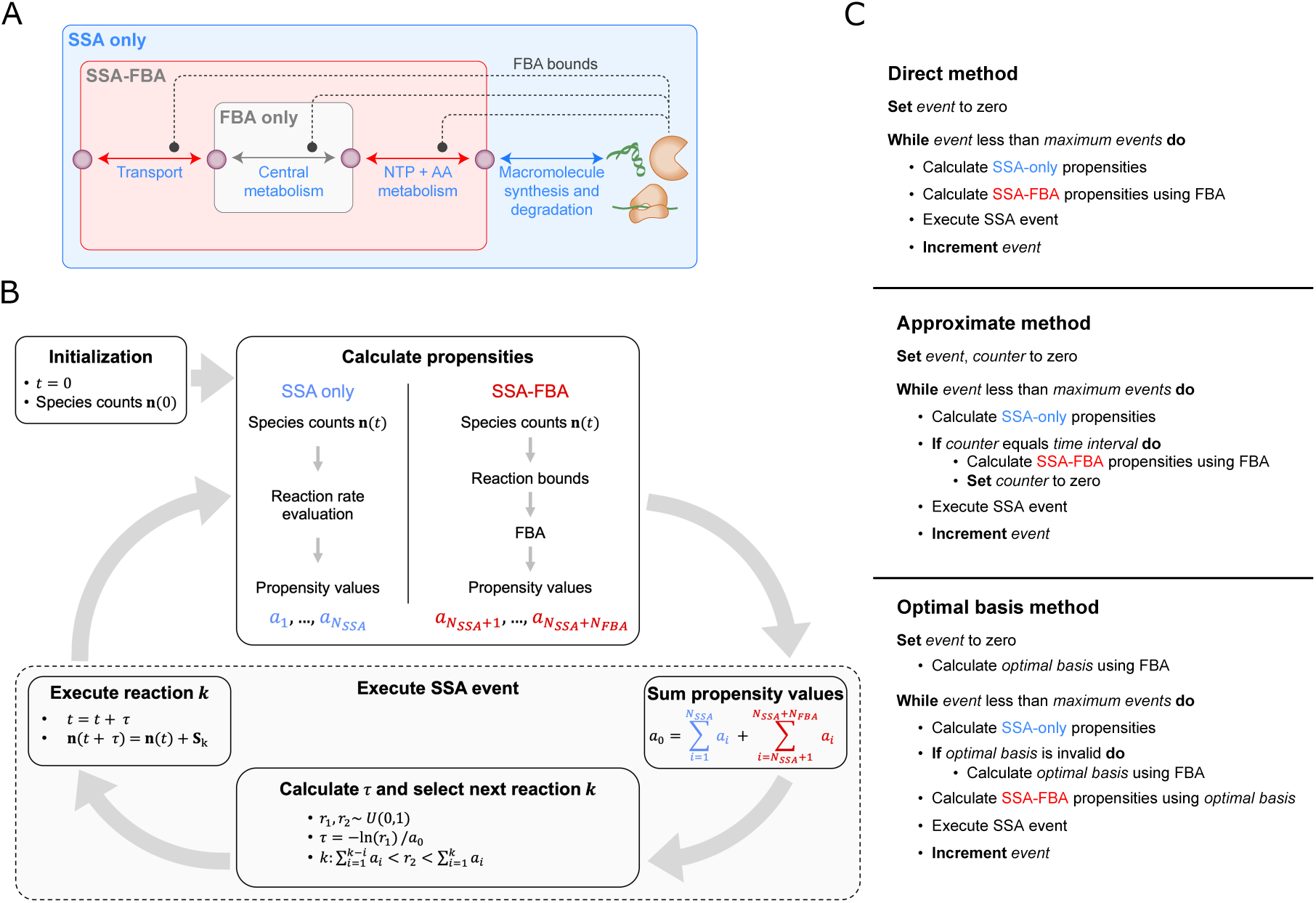
Overview of SSA-FBA model structure and simulation algorithms. **A)** SSA-FBA separates reactions into SSA only (blue), SSA-FBA (red) and FBA only (grey) subsets based on the species that participate in each reaction. In this example, which models the coupling of transmembrane transport and central metabolism to macromolecule synthesis and degradation via amino acid (AA) and nucleotide triphosphate (NTP) metabolism, metabolites in central carbon metabolism are internal to the metabolic reaction network. Gene products such as enzymes and transporters contribute to the rates or propensity values of FBA and SSA-FBA reactions, and this is modelled through FBA bounds on these reactions (dotted lines). **B)** Following initialisation, each iteration of an SSA-FBA simulation first calculates the propensity values for SSA only reactions (blue) by direct evaluation of reaction rate functions (as in Gillespie’s original algorithm, (30, 31)) and calculates optimal propensity values for SSA-FBA reactions (red) using an optimal solution of the FBA model, where current species counts are used in reaction bounds. Next, both types of propensity values are combined to select the next reaction and its execution time. Finally, species counts are updated according to the stoichiometry of the selected reaction. **C)** Pseudo code for the direct, approximate and optimal basis SSA-FBA simulation methods. The approximate method speeds up SSA-FBA simulation by only calculating SSA-FBA propensity values at a fixed time interval (represented by Δ_*event*_ in main text). The optimal basis method speeds up SSA-FBA without approximation by only calculating SSA-FBA propensity values when the optimal basis has changed.

- **FBA only reactions**: reactions that interconvert among the internal species,
- **SSA only reactions**: reactions that interconvert among the external species,
- **SSA-FBA reactions**: reactions that convert between the internal and external species.

For example, the SSA only reactions may correspond to the synthesis and degradation of gene products, while the FBA only and SSA-FBA reactions may include reactions involved in transmembrane transport and central metabolism.

Since external species, such as enzymes and transporters, often influence the rates or propensity values of metabolic reactions, the FBA only and SSA-FBA reactions can be constrained or bounded by the dynamic counts of external species (Figure 2A). Mathematically, this can be expressed as the following linear programming (LP) problem

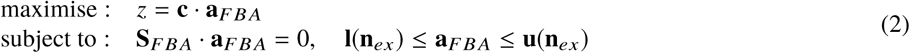

where **a**_*FBA*_ is a vector containing the propensity values of FBA only and SSA-FBA reactions, **S**_*FBA*_ is the sub-matrix of **S** encoding the stoichiometry of the metabolic reaction network, and **l**(**n**_*ex*_), **u**(**n**_*ex*_) are bounds that depend on the counts of external species, **n**_*ex*_ (see Supplementary Appendix S1 for extended discussion). The functional forms of bounds **l**(**n**_*ex*_), **u**(**n**_*ex*_) rely on the way that different reactions are represented in the model: for example, bounds for transport reactions could depend on species counts analogously to the way that exchange flux bounds can depend on extracellular substrate concentrations in DFBA (32), while propensity values of enzymatic reactions can be bounded in proportion to the abundances of intracellular enzyme molecules (so-called enzyme capacity constraints, e.g. (20, 40)). These considerations were implemented in the whole-cell model described in (28) and also the reduced model of *M. pneumoniae* presented as an SSA-FBA case study in this paper (see Supplementary Appendix S3). The coefficient vector **c** is chosen to reflect a biologically-relevant objective such as maximising the rate of production of the metabolites required for growth. Solving the LP problem (2) for a given instance of the external species counts vector **n**_*ex*_ returns an optimal set of propensity values for FBA only and SSA-FBA reactions, while propensity values for the SSA only reactions are calculated from **n**_*ex*_ by direct evaluation of a reaction rate function as in SSA (30, 31).

Next, as outlined in Figure 2B, SSA-FBA combines propensity values for *N*_*FBA*_ SSA-FBA reactions obtained from an optimal solution to the LP problem (2) with the propensity values of *N*_*SSA*_ SSA only reactions to determine the next reaction event according to Gillespie’s original algorithm (30, 31) (although this step can also be replaced by more advanced methods such as the Next Reaction Method (41) or possibly incorporated into tau-leaping-based approximation algorithms (42)). Since FBA only reactions do not affect the counts of external species, their propensity values are not required at this stage. Finally, execution of either an SSA only or SSA-FBA reaction updates the species counts vector, which is in turn used to update the bounds of both FBA only and SSA-FBA propensity values in (2). It is important to highlight that SSA-FBA is not restricted to cases where the metabolic portion of a model is represented by an LP problem, as framed here, but can be extended more generally to scenarios where non-linear optimisation problems used to calculate SSA-FBA propensity values are embedded within SSA. For example, including thermodynamic constraints can further restrict the possible set of SSA-FBA propensity values and results in a non-linear or mixed-integer LP problem (e.g. (43)); however, the efficient simulation algorithm presented in the next section can not necessarily simulate such SSA-FBA models because it depends on the optimal basis structure of LP problems.

In summary, SSA-FBA embeds FBA within SSA by using an LP problem to predict an optimal propensity value for each metabolic reaction, using the counts of species predicted by SSA. This bidirectional coupling of SSA and FBA is analogous to the coupling of FBA and ODE integration in DFBA (32). One challenge of both DFBA and SSA-FBA is that solutions to (2)– and therefore the optimal propensity values of SSA-FBA reactions– are not necessarily unique (44). However, it turns out to always be possible to formulate a lexicographic version of the LP problem by including multiple biological objective functions to be optimised sequentially in order to guarantee uniqueness and retain compatibility with the implementation described in the next section (see also (45)), although construction of a lexicographic LP is, in general, not unique either and imposes additional biological assumptions. It is important to highlight that, unlike variants of SSA that approximate trajectories of the CME (1) in order to reduce the computational overhead (36–39), the goal of SSA-FBA is only to approximately predict the metabolic dynamics of single cells in the absence of detailed kinetic information about each species and reaction (16, 17). The modeller can control the degree of this approximation by how they choose to partition reactions into the three subsets (FBA only, SSA only, or SSA-FBA). As a rule of thumb, we recommend that modellers distinguish between internal and external species (and hence partitioning of reactions) based loosely on the time-scales of the species, but often the distinction between SSA-FBA and FBA only reactions will be largely determined by the structure of the metabolic model and the metabolites that are considered substrates for the gene expression (SSA only) reactions. Further guidelines for encoding SSA-FBA models and additional information about the relationship between SSA-FBA and the CME are outlined in Supplementary Appendix S1.

## Efficient implementation of SSA-FBA

As described above, SSA-FBA iteratively (i) uses the counts of the species and reaction rate laws to calculate the propensity values for each SSA only reaction, (ii) uses the counts of the species in FBA to calculate the propensity values of each SSA-FBA reaction, (iii) uses SSA to select the next reaction and (iv) updates the species counts based on stoichiometry of the selected SSA only or SSA-FBA reaction. The direct method for performing an SSA-FBA simulation in this way is summarised in the top row of Figure 2C. At first glance, the direct method appears to be computationally expensive for larger models because it appears to require solving one FBA problem per SSA execution event.

One way to reduce the computational cost associated with the direct SSA-FBA simulation method is to approximate the SSA-FBA propensity values by updating them less frequently, assuming a time interval (Δ_*event*_) across which an optimal solution to the embedded FBA problem does not change appreciably. When the end of the interval is reached, the constraints of the embedded FBA problem can be updated using the current counts of species and re-solved for a new set of SSA-FBA propensity values used to parameterise integration over the next interval. This approximate SSA-FBA simulation method (centre row in Figure 2C) is analogous to the simplest implementation of DFBA (32) that has been used in the simulation of whole-cell models (28, 29). To evaluate the performance of the direct and approximate simulation methods, we designed a toy model that contained only two SSA only reactions (R0 and R1), one SSA-FBA reaction (R2, growth rate of *Mycoplasma genitalium*) and one variable FBA bound (on the oxygen transport reaction). The toy model contained three species *S*_0_, *S*_1_, *S*_2_ and is represented by the reaction schema

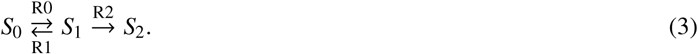

In this schema, the propensity value of the SSA-FBA reaction R2 is the current optimal value of the growth rate objective function calculated by FBA using the metabolic network model of *M. genitalium* from (28). The R2 propensity value thus depends indirectly on species *S*_1_, which bounds the maximal rate of the oxygen uptake reaction in the *M. genitalium* metabolic network model based on the functional form *u*(*S*_1_) = *S*_1_/10. The optimal value of the objective function of the *M. genitalium* metabolic network model obtained by solving the associated FBA problem at this substrate level (corresponding to a given maximal rate of oxygen uptake) is then used as propensity value for reaction R2.

We began simulations with initial species counts *S*_0_ = 1000, *S*_1_ = 0, *S*_2_ = 0, and simulated the toy model across 3000 reaction execution events using either the direct or approximate SSA-FBA simulation method. In particular, we used the toy model to explore how the choice of Δ_*event*_ affects results of SSA-FBA simulations using the approximate method. When Δ_*event*_ is small, the variances of approximate SSA-FBA simulations are comparable to those obtained using the direct method (Figures 3A and 3B) and increasing Δ_*event*_ increases this variance (Figure 3C). The toy model highlighted a shortcoming of the approximate SSA-FBA simulation method that trajectories occasionally (particularly for larger Δ_*event*_ as visible in Figure 3C, but observed for all Δ_*event*_ tested) display numerical instability and diverge from realistic values towards the latter end of a simulation. This type of numerical instability also arises during direct integration of many DFBA models (45), but is particularly problematic for SSA-FBA because its stochastic nature means the pathology cannot be conclusively ruled-out on the basis of trial simulations (for example, to establish optimal size of Δ_*event*_). The origin of the problem is that the embedded FBA problem can become infeasible within an integration interval, which induces a closed domain of definition for the dynamic system (45).

**Figure 3:**
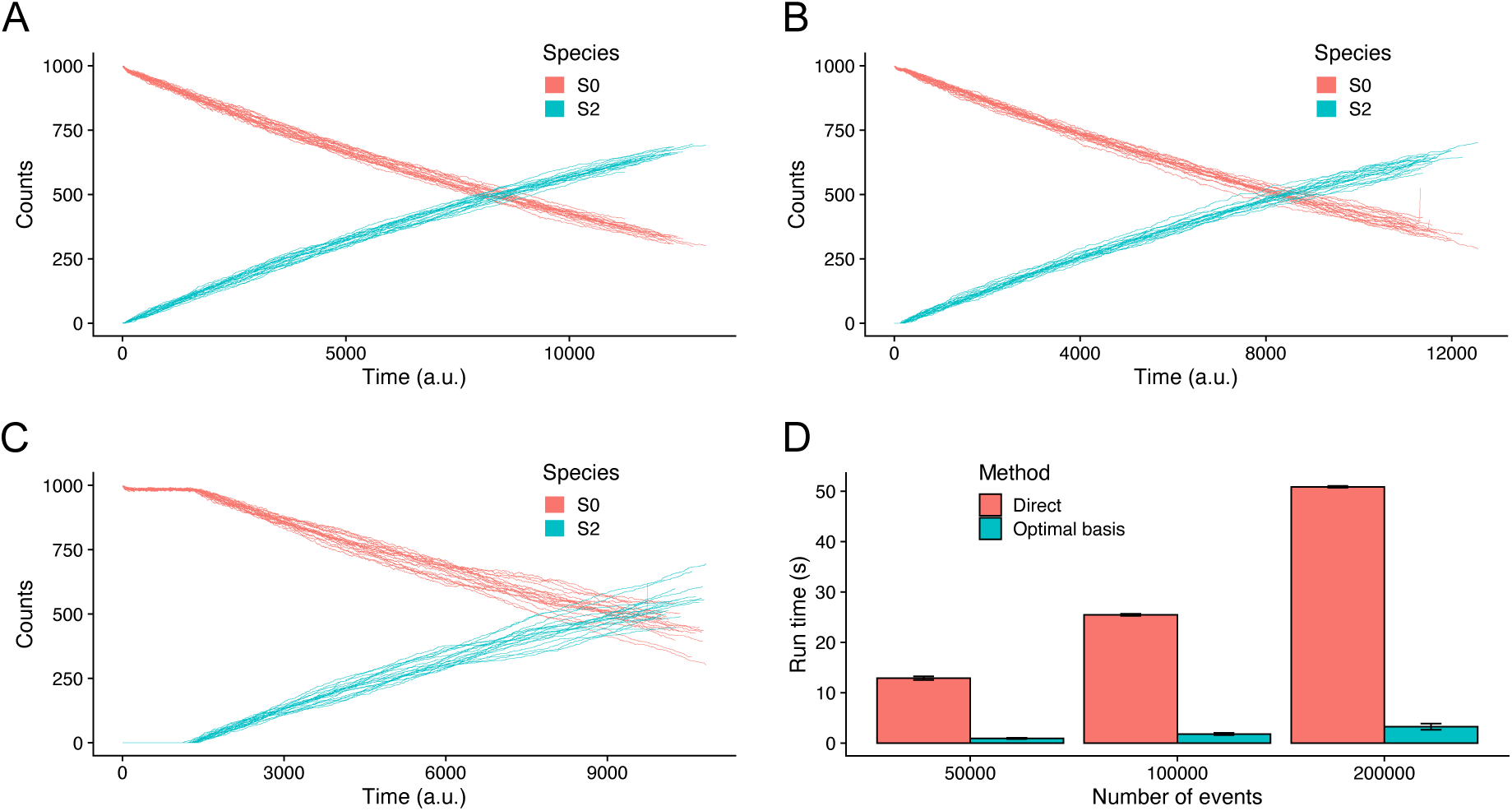
Comparison of the SSA-FBA simulation methods. Trajectories from three sets of 20 representative simulations, each consisting of 3000 reaction execution events: **A)** direct SSA-FBA simulation of toy model (equivalent to Δ_*event*_ = 0); **B)** approximate SSA-FBA simulation of toy model with Δ_*event*_ = 50; **C)** approximate SSA-FBA simulation of toy model with Δ_*event*_ = 500. **D)** Comparison of run times for direct and optimal basis simulation methods. Error bars are standard deviations for run times across four replicate simulations involving 50000, 100000, or 200000 FBA bound updating events (described in Supplementary Appendix S2).

We therefore searched for a strategy to reduce the computational cost of SSA-FBA simulation without the loss of accuracy and numerical stability associated with approximation. Motivated by a recent method for numerical integration of ODEs with embedded LP problems (45), we developed a more efficient algorithm that leverages the fact that FBA problems only need to be resolved when their optimal basis changes (outlined in bottom row of Figure 2C and full details in Supplementary Appendix S2), which typically occurs much less frequently than after the execution of every SSA reaction because of the different timescales of metabolic and gene expression reactions. In this approach, only the validity of the current optimal basis being used to calculate optimal propensity values for SSA-FBA reactions must be established after each reaction execution event, which in the context of most SSA-FBA models is computationally cheaper than solving the embedded LP problem each time molecular species counts are updated. The result of the optimal basis simulation method remains mathematically equivalent to that of the direct SSA-FBA simulation method, but is obtained substantially faster because it requires solving many fewer LP problems. Figure 3D displays representative results of the comparisons (described in Supplementary Appendix S2) between the optimal basis and direct (equivalent to the approximate with Δ_*event*_ = 0) SSA-FBA simulation methods, showing that the former improves run time by an order of magnitude over the latter. Intuitively, the performance of the optimal basis algorithm relative to the direct method will depend on the level of connectivity between the embedded FBA problem and remaining SSA component of a model. Greatest performance increases will be seen in models where most reaction execution events do not require updating many FBA bounds, so that the computational cost associated with validating the current optimal basis is substantially less than that associated with solving the embedded LP problem. Given the typical ways of representing biological coupling between metabolic and macromolecular portions of models (e.g., enzyme capacity constraints where most enzymes are involved in the catalysis of a single reaction (20, 40)), it follows that the connectivity of biologically-realistic reaction networks implies the optimal basis algorithm will significantly improve simulation performance for the vast majority of cases.

### Case study: reduced single-cell model of *M. pneumoniae*

In this section, we present results from a case study using SSA-FBA to simulate the dynamics of metabolism in a reduced model of a single *M. pneumoniae* cell in order to understand how variability in single-cell metabolism is driven by the intrinsic stochasticity of gene expression. *Mycoplasma* have the smallest genomes among known freely living cells and, by the law of large numbers, their small sizes therefore mean they should exhibit the largest effects of stochasticity in the absence of regulation: the relative absence of genetically-encoded regulation compared with higher organisms means that stochastic effects are likely to play a larger role in their metabolic behaviour. Furthermore, *M. pneumoniae* is one of the best-characterised members of this family (46), which makes it an excellent case study for understanding the behavioural effects of stochasticity in single-cell metabolism. Its close relative *M. genitalium* was previously used to build a whole-cell model (28), to which SSA-FBA could also be applied in principle along with genome-scale metabolic reaction networks from other organisms. The purpose of this section is not to study an entire whole-cell model however, as the complexity involved in doing so would extend far beyond the main scope of this paper, whose goal is to introduce SSA-FBA as a modelling framework and demonstrate its feasibility by simulating a model of reasonable size and biological accuracy.

We constructed a model of *M. pneumoniae* consisting of 505 biochemical reactions that account for the physiology of metabolism and the intrinstic drivers of stochasticity in gene transcription, translation and macromolecular degradation. Most parameter values were derived from metabolomics, transcriptomics, and proteomics data about *M. pneumoniae* (47–50), with the exception of rate constants for metabolic reactions (for which many fewer experimental observations are available) that are drawn from a variety of other bacteria (52). The model contains a metabolic reaction network with 86 metabolic reactions (Figure 4A), including a biosynthetic pseudo-reaction that constitutes the objective function of the embedded FBA problem. The biosynthetic pseudo-reaction was constructed to be internally consistent with the SSA only reactions that model gene expression as described below and does not account for additional maintenance energy costs not included the model– a complete description of the biosynthetic pseudo-reaction is provided in the relevant subsection of Supplementary Appendix S3. There are 81 protein species that either function individually or in complex (27 complexes in total) as enzymes or transporters regulating flux through the FBA reactions, or serve a direct role in gene expression (ribosomal proteins, RNA polymerase or RNAse subunits). Furthermore, each protein species is associated with an mRNA molecule (in addition to the three ribosomal RNAs) and four reactions corresponding to RNA transcription, RNA degradation, protein synthesis and protein degradation, which together with complexation reactions makes 420 SSA only reactions.

**Figure 4:**
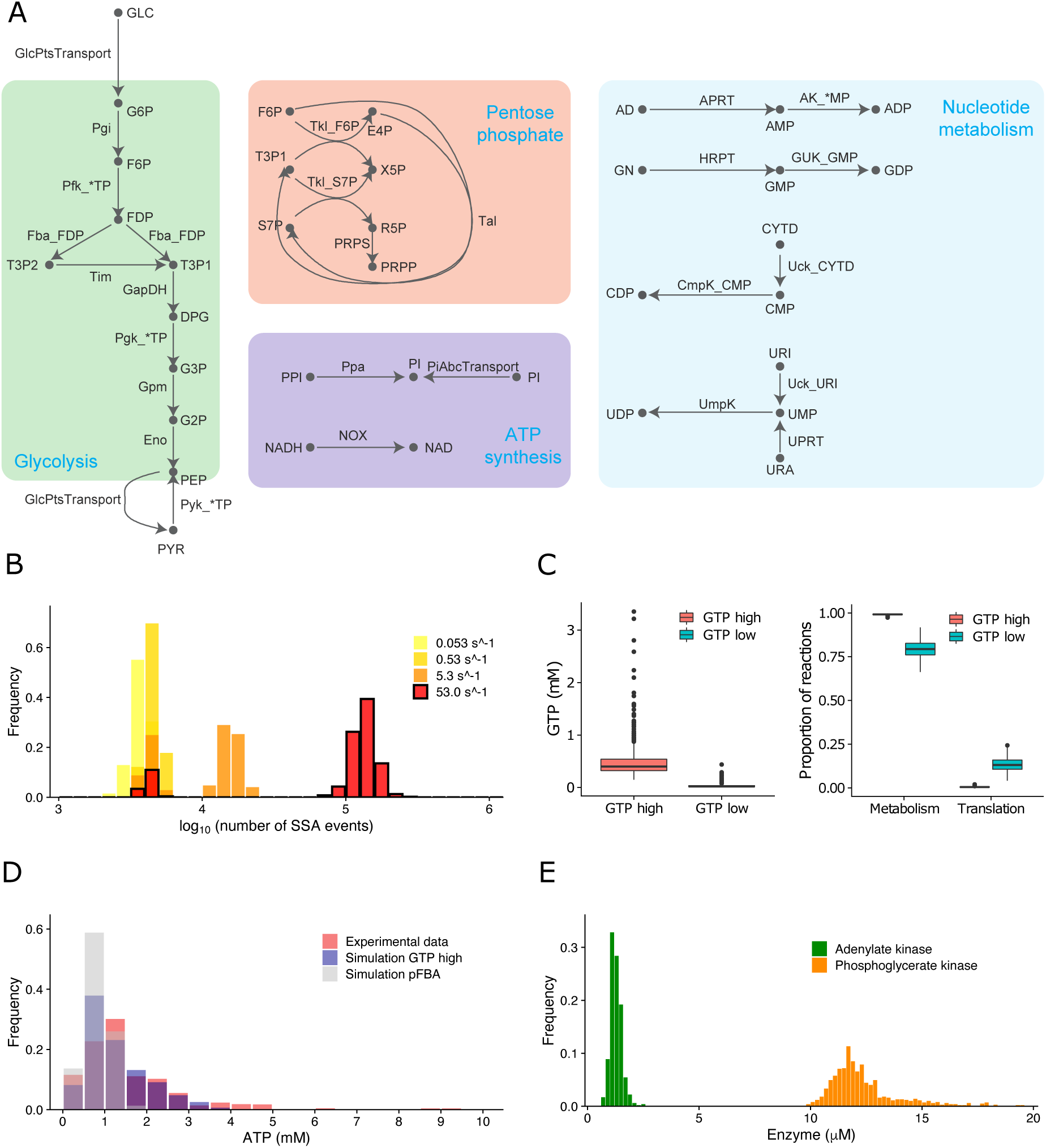
Single-cell simulation of *M. pneumoniae* metabolism using SSA-FBA. **A)** Metabolic reaction network used in the model annotated with species listed in Supplementary File S1. **B)** Comparison of distributions of number of SSA execution events per simulation for different values of the GMK reaction rate constant. For reference, the bimodal distribution corresponding to *k*_*cat*_ = 53.0 *s*^−1^ is outlined in black. **C)** Box plots showing significant differences between GTP concentrations (*p* < 2.2 × 10^−16^), and the relative proportions of reaction execution events that correspond to metabolic reactions (*p* < 2.2 × 10^−16^) and translation reactions (*p* < 2.2 × 10^−16^), respectively, between the low and high GTP groups. **D)** Distributions of time-averaged ATP concentrations from simulations with GTP high group and pFBA compared with the experimental distribution from (54). **E)** Distribution of time-averaged ADK and PGK concentrations from simulations with GTP high group.

As described previously, it is natural that metabolite species (NTPs and AAs) appearing as reactants in the rate laws for the SSA only reactions governing macromolecular synthesis and degradation are viewed as external species of the metabolic reaction network along with extracellular substrates, while all other metabolites are considered internal. Therefore, reactions of the embedded FBA model involved in the production or consumption of NTPs and AAs or extracellular transport are naturally assigned to the SSA-FBA reaction subset. The remaining reactions of the embedded FBA model (those whose substrates or products do not involve NTPs, AAs or extracellular substrates) are consequently assigned to the FBA only subset. Supplementary File S1 contains all of the compartments, species, reactions, rate laws, and parameters; Supplementary Appendix S3 provides an extended description of the model.

The reduced single-cell model is based partly on metabolic rate constants measured for bacteria distantly related to *M. pneumoniae* and so should not be considered a precise description of *Mycoplasma* physiology. Indeed, even the most detailed constraint-based, genome-scale models attempting to provide descriptions of entire cell physiology (51) must contend with the problem of missing parameter values, and the dependance of their completeness on future experimental research is something that genome-scale SSA-FBA simulations are subject to also. The average fold variation for wild-type bacterial enzymatic reaction rate constants (*k*_*cat*_ values) in BRENDA (52) is 3951.6, as measured by 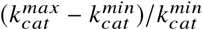 (where 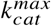 and 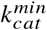 are the largest and smallest, respectively, *k*_*cat*_ values reported for that enzyme) and averaged across all enzymes included in the model, suggesting one or more of these model parameters require calibration in order to at the very least generate predictions consistent with the observed growth physiology of *M. pneumoniae*.

Here one advantage coming from computational efficiency of the optimal basis SSA-FBA simulation algorithm is making it possible to parsimoniously search metabolic parameter space and establish parameter sensitivities by individually varying enzymatic reaction rate constants through repeated simulation. As an illustrative example, we focus on the sensitivity of growth rate to variations in the *k*_*cat*_ value of guanylate kinase (GMK, gene MPN246) as a key enzyme (a phosphotransferase) forming part of the phosphotransfer network responsible for the homeostasis of NTPs and cellular energetics (53). To generate each set of simulation results for a given GMK *k*_*cat*_ value we ran a thousand instances of the single-cell *M. pneumoniae* model using the optimal basis SSA-FBA method initialised with counts of species randomly sampled from a Poisson distribution parametrised by their mean values across the cell cycle. In each case we analysed simulations over a 25 min interval of biological time, across which cell volume is assumed to be approximately constant given the relatively long (6-8 hrs) doubling time of *M. pneumoniae* (46), excluding the first 25 mins of each 50 min simulation to prevent initial transients from contaminating the analysis.

The *k*_*cat*_ value of GMK was varied from its lowest measured value (equal to the *k*_*cat*_ of adenylate kinase (ADK) in *Bacillus subtilis*, a much faster growing gram-positive bacteria), by increasing in 10-fold increments towards its highest estimated value (ADK in *Vibrio natriegens*). Notably, we found that the distribution of SSA execution events per simulation becomes bimodal as the GMK reaction rate constant increases (Figure 4B). This bimodal distribution at larger GMK *k*_*cat*_ values computationally separates simulations into two distinct groups with a low (less than 10,000 SSA execution events) or high (more than 10,000 SSA execution events) number of SSA execution events per simulation, which are biologically associated with significantly low or high time-averaged GTP concentrations: 0.04 ± 0.06 mM vs 0.47 ± 0.29 mM (mean ± SD) in low vs high group, respectively (Figure 4C). Figure 4C also shows that, for the largest value of the GMK reaction rate constant tested, simulations in the low GTP group (146 out of 1000 simulations) were associated with a significantly lower average proportion of SSA execution events corresponding to metabolic reactions compared to simulations in the high GTP group: 79.36 % vs 99.16 % in low GTP group vs high GTP group, respectively, implying that higher time-averaged GTP concentrations are associated with increased metabolic activity. Conversely, a significantly relatively higher average proportion of SSA execution events correspond to translation reactions in the low GTP group compared to the high GTP group: 13.46 % compared with 0.50 % in low GTP group vs high GTP group, respectively, consistent with the fact that translation is a major consumer of intracellular GTP. Given the critical importance of ATP as source of energy for intracellular reactions supporting growth, we then studied the distribution of time-averaged ATP concentrations across the population of simulated cells. We found that the time-averaged concentrations ATP across the group of metabolically more active cells is positively skewed in qualitative agreement with experimental measurements of ATP concentrations inside individual bacteria (54) (Figure 4D): 1.28 ± 0.75 mM vs 1.54 ± 1.22 mM (mean ± SD) and skew values of 1.16 vs 2.20 for simulation vs experiment, respectively. Conversely, the distribution of ATP concentrations in metabolically less active cells was slightly negatively skewed: 0.81 ± 0.22 mM (mean ± SD) and skew value of −0.35.

To understand whether multiple optimal solutions of the embedded FBA problem explain the bimodal simulation behaviour observed at larger GMK *k*_*cat*_ values, we performed flux variation analysis (FVA) (44) with the metabolic reaction network at steady state. FVA revealed that the reactions catalysed by GMK, ADK and phosphoglycerate kinase (PGK; ATP and GTP production) are able to carry arbitrarily large flux values within the optimal solution space defined by maximising flux through the biosynthetic pseudo-reaction. Correspondingly, elementary flux mode enumeration (55) used to identify all minimal pathways through the metabolic reaction network revealed a single internal cycle (and its reverse) that contains the support of the GMK, ADK and PGK reactions without net metabolite production or consumption. Such internal cycles, not involving the primary exchange reactions representing exchange of material between the model and the environment, are known to violate the first law of thermodynamics in conventional constraint-based models (43), but in the reduced *M. pneumoniae* model GMK,

ADK and PGK catalyse SSA-FBA reactions involved in internal exchange of NTPs. We confirmed that simulations within the group of metabolically more active cells have significantly higher mean numbers of these four reactions executing across the course of a simulation than any other SSA-FBA reaction (*p* < 2.2 × 10^−16^, based on one sample t-test between the mean number of execution times of a given SSA-FBA reaction compared to that of all others), which is a result of the internal loop being able to carry a larger flux value in the optimal solution space when the GMK reaction rate is bounded by a higher *k*_*cat*_ value. We also found a significant positive correlation between time-averaged ATP concentrations and ADK but not PGK enzyme levels (*R* = 0.18, *p* < 10^−7^ and *R* = 0.04, *p* = 0.24, respectively, based on Pearson’s product moment correlation coefficient), which were positively skewed (1.24 ± 0.26 μM and 12.35 ± 1.52 μM (mean ± SD) and skew values 1.00 and 1.67, respectively (Figure 4E), suggesting that asymmetric metabolite distributions are perhaps the result of fluctuating ADK levels controlling the relative levels of flux through the internal loop. Conversely, in the group of metabolically less active cells, reactions catalysed by GTP phosphofructokinase (PFK), PGK, and GTP pyruvate kinase (PYK) along with the glucose import reaction are significantly overrepresented in simulations (*p* < 2.2 × 10^−16^), without significant correlation between between time-averaged ATP concentrations and ADK enzyme levels.

Since the internal cycle involving ADK responsible for positively skewed time-averaged ATP concentrations is deemed thermodynamically infeasible by constraint-based modelling criteria (43), we next employed lexicographic optimisation with the biosynthetic pseudo-reaction and a second parsimonious FBA objective (pFBA: minimising the sum of absolute flux values, see (56) and Supplementary Appendix S3). Although imposing a strong assumption on the biology of *M. pneumoniae* metabolism, pFBA is guaranteed to return a set of optimal SSA-FBA propensity values that are calculated without the involvement of internal cycles. Subsequently, the distribution of SSA execution events per simulation with pFBA retained a single mode even at the highest value of the GMK reaction rate constant and SSA-FBA execution events associated with ATP PFK, PGK, ATP PYK and glucose import were significantly overrepresented (*p* < 2.2 × 10^−16^). Mean ATP concentrations with the pFBA objective were closer to those of previously metabolically less active cells, although the distribution retained a slight positive skew: 0.83 ± 0.01 mM (mean ± SD) and skew value of 0.30 (Figure 4D). Taken together, the requirement to include thermodynamically infeasible internal cycles to reproduce experimentally observed ATP distributions is a critical shortcoming of the reduced *M. pneumoniae* single-cell model that emerges because SSA-FBA reactions involved in internal NTP exchange are coupled to macromolecular synthesis and degradation. This issue would not arise in typical population-based DFBA models where the embedded FBA problem is only coupled to the ODE via its primary exchange reactions. It implies that additional non-growth-associated NTP maintenance reactions should be incorporated into single-cell models to account for alternative NTP requirements (57) and more formally mediate internal NTP exchange, possibly with additional constraints that impose conservation on certain metabolite pools (58).

## DISCUSSION

In this paper we have presented SSA-FBA, the first framework with a realistic potential for simulating the complete dynamics of metabolic networks in single cells at the resolution of individual species and reactions. SSA-FBA is a hybrid method for embedding FBA in SSA that is well-suited to simulating metabolic networks for which detailed kinetic information is often lacking. We also have developed an advanced optimal basis algorithm for efficiently executing SSA-FBA without approximation. A case study using our algorithm to simulate a reduced model for the metabolism of an individual *M. pneumoniae* cell demonstrates that SSA-FBA has the potential to reveal how stochasticity at the single-cell level contributes to metabolic heterogeneity at the population level. Our results may help to identify the sources of variation observed in experimental measurements of important metabolites in single cells, indicating that reaction network coupling could contribute to population heterogeneity by amplifying or attenuating the sources noise in an integrative fashion. SSA-FBA can therefore be used to probe metabolic heterogeneity in a manner complementary to alternative existing experimental and computational methods (11, 22, 25, 27).

Limitations of the current study include various assumptions involved in the separation of timescales between reactions from metabolic portions of models and reactions responsible for macromolecular synthesis and degradation. In particular, although the faster timescales of metabolic reactions and higher abundance levels of metabolites have been widely used to justify a reduction of the CME or deterministic descriptions of metabolism (25, 37–39), the validity of these assumptions should be questioned for each individual representation of single-cell metabolism prior to implementation of SSA-FBA. We have also highlighted two additional restrictions on SSA-FBA that reflect general limitations of any metabolic modelling framework based on an LP formulation, but can be overcome using extensions of the work presented here: first, uniqueness of SSA-FBA propensity values is not guaranteed, although this can be achieved by implementing lexicographic optimisation compatible with the efficient SSA-FBA simulation algorithm; secondly, whilst not possible to simulate using the efficient algorithm, SSA-FBA models can be generalised to include scenarios where a non-linear optimisation problem is used to represent metabolism. Finally, the general lack of experimentally-determined parameter values that plagues current whole-cell modelling attempts (28, 29, 51) serves as a current limitation to building large-scale, single-cell models of metabolism, but, as further advancements are made in data collection, SSA-FBA will retain an advantage over alternative frameworks that depend on precise knowledge of kinetic parameters.

Future extensions of our work will involve applying SSA-FBA to larger, more realistic models of entire cells such as those based on resource balance analysis (51), in addition to reconsidering how standard constraint-based formulations of metabolite pools, energy maintenance and objective functions should be adapted to suit the biological nature of single-cell biology. More complex models of single-cell metabolism could include mechanisms that impart regulatory control of stochasticity (59), and pave the way for combining insights from simulation with experimental advances in microfluidics (60) or real-time quantification of RNA translation events within individual cells (61). From an algorithmic perspective, SSA-FBA could be extended to incorporate additional time-scales governed by continuous stochastic or deterministic processes (e.g., (62)), and its computational efficiency perhaps further enhanced through parallelisation methods (63).

## Supporting information

Supplementary model file

## AUTHOR CONTRIBUTIONS

DST designed research and implemented algorithms and software with conceptual feedback from other coauthors. JRK designed case study and DST implemented and performed simulations. DST wrote the article with input from all authors.

## DECLARATION OF INTERESTS

The authors declare no competing interests.

## ACKNOWLEDGMENTS

Part of this work was completed while DST was a Simons Foundation Fellow of the Life Sciences Research Foundation hosted by Columbia University Irving Medical Center. The work of JRK was supported by National Science Foundation award 1649014 and National Institutes of Health award R35GM119771. We thank ME Beber and J Carrasco Muriel for their close collaboration with the first author on a related project that provided some of the motivation for the current study, and Maria Lluch-Senar, Veronica Llorens, and Luis Serrano for discussions on the physiology of Mycoplasmas. The authors declare that they have no conflict of interest.

## SUPPLEMENTARY MATERIAL

An online supplement to this article can be found by visiting BJ Online at http://www.biophysj.org.

## Notes

https://gitlab.com/davidtourigny/single-cell-fba

### Competing Interest Statement

The authors have declared no competing interest.

### Summary of Updates

Final accepted version accommodating referee comments

