## Supplementary appendix for "Simulating single-cell metabolism using a stochastic flux-balance analysis algorithm"

October 1, 2021

#### S1. Extended description of SSA-FBA

In this section of the Appendix we expand upon the relationship between SSA-FBA and the reduced chemical master equation. Reduction of the chemical master equation (CME) is based on an assumption of time-scale separation, i.e., the assumption that transients in concentrations of a group of $N^s$ (slow) species decay much slower than those of the remaining group of $N^f$ (fast) species. The vector of total molecular counts of $N = N^f + N^s$ species, $\mathbf{n} = (\mathbf{n}^f, \mathbf{n}^s)$, is then partitioned into fast $\mathbf{n}^f$ and slow $\mathbf{n}^s$ vectors of dimensions $N^f$ and $N^s$, respectively. In SSA-FBA, partitioning of molecular species is based on choices made by the modeller when designating species to be internal or external to the metabolic network, which in turn dictates the structure of the SSA-FBA model to be simulated. Ideally, we would like to enforce that internal species correspond to fast species and external to slow, but sometimes the formal derivation of SSA-FBA described below may break down because the identification of $\mathbf{n}^f$ with counts of internal species and $\mathbf{n}^s$ with counts of external species is not realised by a model. It should be pointed out that an equivalent limitation holds for dynamic FBA (DFBA) [1], where conventionally the distinction between fast and slow concentrations is based on the separation of intra- and extracellular metabolites.

Given $M$ chemical reactions with corresponding propensity values $a_j$ ($j = 1, 2, ..., M$), the full CME [2] expressed in terms of the partitioning of species is

$$\frac{dP(\mathbf{n}^f, \mathbf{n}^s)}{dt} = \sum_{j=1}^{M} a_j(\mathbf{n}^f - \mathbf{S}_j^f, \mathbf{n}^s - \mathbf{S}_j^s) P(\mathbf{n}^f - \mathbf{S}_j^f, \mathbf{n}^s - \mathbf{S}_j^s) - a_j(\mathbf{n}^f, \mathbf{n}^s) P(\mathbf{n}^f, \mathbf{n}^s) \tag{1}$$

where $\mathbf{S}_j^f$ and $\mathbf{S}_j^s$ are the $j$th columns of the portions of the stoichiometric matrix corresponding to fast and slow species, respectively. Summing over $\mathbf{n}^f$ we obtain a master equation for the marginal distribution

$$\frac{dP(\mathbf{n}^s)}{dt} = \sum_{j=1}^{M} \bar{a}_j(\mathbf{n}^s - \mathbf{S}_j^s) P(\mathbf{n}^s - \mathbf{S}_j^s) - \bar{a}_j(\mathbf{n}^s) P(\mathbf{n}^s), \tag{2}$$

where

$$\bar{a}_j(\mathbf{n}^s) \equiv E\left[a_j(\mathbf{n}^f, \mathbf{n}^s) | \mathbf{n}^s\right] = \sum_{\mathbf{n}^f} a_j(\mathbf{n}^f, \mathbf{n}^s) P(\mathbf{n}^f | \mathbf{n}^s) \tag{3}$$

is the expectation value of $a_j(\mathbf{n}^f, \mathbf{n}^s)$ conditioned on $\mathbf{n}^s$. We will refer to Equation (2) as the marginal chemical master equation (MCME). Rao and Arkin [3] introduced a stochastic equivalent of the quasi-steady state assumption (sQSSA): that the conditional distribution $P(\mathbf{n}^f | \mathbf{n}^s)$ is assumed to rapidly converge to a stationary distribution on an interval over which $P(\mathbf{n}^s)$ remains approximately constant

$$0 \approx \frac{dP(\mathbf{n}^f | \mathbf{n}^s)}{dt} = \left( \frac{dP(\mathbf{n}^f, \mathbf{n}^s)}{dt} - P(\mathbf{n}^f | \mathbf{n}^s) \frac{dP(\mathbf{n}^s)}{dt} \right) \frac{1}{P(\mathbf{n}^s)}. \tag{4}$$

After substituting $dP(\mathbf{n}^f, \mathbf{n}^s)/dt$ and $dP(\mathbf{n}^s)/dt$ for their right-hand sides in Equation (4), this condition together with the MCME (2) yields a differential algebraic equation (DAE) for $P(\mathbf{n}^s)$. Alternatively, the work of Thomas, Straube & Grima [4, 5] uses a projection operator method combined with linear noise approximation to obtain a reduced Fokker-Planck equation for the fluctuations in the concentrations of the slow variables, and Smith, Cianci & Grima [6] present a different reduction of the CME based on the separation of chemical abundances (e.g., metabolites considered abundant and macromolecules considered non-abundant) rather than time-scales.

Full reduction of the CME (1) depends on obtaining the stationary conditional distribution $P(\mathbf{n}^f|\mathbf{n}^s)$, which is required for calculating each $\bar{a}_j(\mathbf{n}^s)$ that is then substituted into the MCME (2). Due to analytic intractability of Equation (4) however, a closed form expression for $P(\mathbf{n}^f|\mathbf{n}^s)$ is usually inaccessible and therefore this distribution and $\bar{a}_j(\mathbf{n}^s)$ must be approximated for most systems. The approximate form of $P(\mathbf{n}^f|\mathbf{n}^s)$ used in [3, 7] is Markovian, whereas in [4, 5, 8] a number of arguments are presented to justify using a Gaussian distribution built from solutions of the associated deterministic rate equations and covariance matrix from first-order corrections. These deterministic rate equations correspond to the lowest-order terms in the system size expansion described in [4, 5], but can also be obtained by full marginalisation of the CME (1) to obtain ordinary differential equations (ODEs) for expected counts molecular species in terms of the expected propensity values $E\left[a_j(\mathbf{n}^f,\mathbf{n}^s)\right]$ $(j = 1,2,...,M)$. Rate equations are more commonly considered in terms of concentrations of species $\mathbf{x} = \mathbf{n}/\Omega$ and macroscopic rate functions $v_j(\mathbf{x}) = a_j(\mathbf{n})/\Omega$ (where $\Omega$ is the system's volume), which motivates the system size expansion that becomes accurate in the limit $\Omega \to \infty$. In [8] the system size expansion combined with time-scale separation produces an approximation of expected propensity values based on conditional averages for fast species satisfying the deterministic steady state condition, whereas in [3, 7] each $\bar{a}_j(\mathbf{n}^s)$ is constructed using moments of the Markovian approximation for $P(\mathbf{n}^f|\mathbf{n}^s)$.

The key motivation for SSA-FBA is that, in large-scale metabolic network models, rate equations are rarely known (even approximately) for fast reactions, and therefore it becomes impossible to parameterise any exact or approximate form of the MCME described above. Moreover, even if the functional form of $\bar{a}_j(\mathbf{n}^s)$ could be justified on biological grounds, it would almost never be the case that all parameter values on which propensity functions depend could be accurately measured experimentally. Consequently, SSA-FBA borrows from the solution to this problem proposed by

(D)FBA [1, 9, 10] where linear programming (LP) is used to identify a numerical solution to the deterministic algebraic steady state conditions

$$\sum_{j=1}^{M} \mathbf{S}_j^f \cdot \bar{a}_j(\mathbf{n}^s) = 0. \tag{5}$$

Using LP to find numerical values for expected propensities $\bar{a}_j(\mathbf{n}^s)$ satisfying constraints (5) and then substituting these into the MCME (2) will formally define SSA-FBA as a stochastic extension of DFBA. Indeed, marginalising the MCME (2) and expressing the resulting ODEs and constraints (5) in terms of species concentrations yields

$$\frac{d\mathbf{x}^s}{dt} = \sum_{j=1}^{M} \mathbf{S}_j^s \cdot v_j(\mathbf{x}^s), \quad \sum_{j=1}^{M} \mathbf{S}_j^f \cdot v_j(\mathbf{x}^s) = 0, \tag{6}$$

which is precisely the DAE used in the definition of DFBA.

In an SSA-FBA model, reactions are separated into three mutually disjoint subsets based on whether the participating species are defined to be internal or external to the metabolic reaction network (see Section 2 in the main text for illustration). Since to some extent the validity of SSA-FBA is dependent on *internal* species being identified with *fast* species and *external* with *slow*, respectively, these two sets of terms will be used interchangeably in the sections below. As we have already pointed out, to strengthen the formal relationship between SSA-FBA and the reduced CME described above, it is important that this identification is adhered to whenever possible. The three subsets of reactions in an SSA-FBA model are:

- **FBA only reactions**: reactions that interconvert among the internal species; their *bounds* can be calculated using the counts of slow species,
- **SSA only reactions**: reactions that interconvert among the external species,
- **SSA-FBA reactions**: reactions that convert between the internal and external species; their *bounds* can be calculated using the counts of slow species.

SSA-FBA uses FBA to obtain numerical values for the expected propensity values of FBA only and SSA-FBA reactions since both depend on fast species. On the other hand, propensity values for SSA only reactions can be calculated by evaluating the rate law directly because, by definition, these reactions have expected propensity values given by $\bar{a}_j(\mathbf{n}^s) = a_j(\mathbf{n}^s)$. To calculate propensity values for the subset of $M_{FBA}$ (FBA only and SSA-FBA) reactions at time $t$, LP maximises a linear objective function to obtain an $M_{FBA}$-dimensional vector $\bar{\mathbf{a}}_{FBA}$ of FBA only and SSA-FBA propensities that satisfy (5) and fall within lower and upper bounds $\mathbf{l}(t) \equiv \mathbf{l}(\mathbf{n}^s(t))$ and $\mathbf{u}(t) \equiv \mathbf{u}(\mathbf{n}^s(t))$. As the notation suggests, these bounds are parameterised by the current molecular counts $\mathbf{n}^s(t)$ of slow species at time $t$. The embedded LP problem of SSA-FBA at time $t$ is

$$\begin{aligned} \text{maximise}: \quad & z = \mathbf{c} \cdot \bar{\mathbf{a}}_{FBA} \\ \text{subject to}: \quad & \mathbf{S}^f \cdot \bar{\mathbf{a}}_{FBA} = 0, \quad \mathbf{l}(t) \leq \bar{\mathbf{a}}_{FBA} \leq \mathbf{u}(t), \end{aligned} \tag{7}$$

where $\mathbf{S}^f$ is an $(N^f \times M_{FBA})$-dimensional matrix and $\mathbf{c} = (c_1, c_2, ..., c_{M_{FBA}})^T$ an $M_{FBA}$-dimensional vector of constant coefficients. The resulting SSA-FBA propensity values obtained by solving the LP problem (7) are combined with those of the SSA only reactions to determine the next reaction event occurring at time $t+\tau$ as determined by Gillespie's stochastic simulation algorithm (SSA) [11, 12], which would provide an exact trajectory of the MCME (2) if propensity values were known precisely. Following execution of the selected reaction using the corresponding stoichiometry, updated molecular counts $\mathbf{n}^s(t+\tau)$ of slow species are then used to update the bounds in the LP problem, which can be solved for a new set of FBA only and SSA-FBA propensity values. The procedure is repeated until the end of the simulation is reached.

The relative scale of SSA only to SSA-FBA propensity values obtained from the embedded FBA problem (7) serves as an additional parameter of an SSA-FBA model, as are the stoichiometry values of the reactions these correspond to. The choice of this relative scaling factor proves to be important for simulation efficiency and is related to the multi-scale nature of single-cell metabolism, because numerical exploration of the state space will become challenging if propensity values differ by several orders of magnitude. There is some flexibility regarding the relative scaling of SSA only to SSA-FBA propensity values; our provided guidelines are that this scale factor should be inversely correlated with the stoichiometry of SSA-FBA reactions (i.e., stoichiometry of SSA-FBA reactions should increase as their propensity values scale down relative to SSA only propensity values).

### S2. Fast algorithm for exact SSA-FBA simulations

As outlined in Appendix S1, exact simulation of an SSA-FBA model appears to imply that a call to an LP solver (such as the Simplex Method) must be made following execution of every reaction event, which could become computationally expensive for systems where execution of many reaction events is necessary before any appreciable effects of changing the bounds of FBA only or SSA-FBA reactions are realised. However, in this section of the Supplementary Appendix, we detail an advanced optimal basis method which typically requires many fewer LP executions. This significantly improves the run time of an SSA-FBA simulation, while still executing exact simulations in the sense described in Appendix S1. Implementation of this method involves two algorithms: Algorithm 1 is a simple extension of SSA [11, 12] for the MCME (2) that incorporates propensity values of SSA-FBA reactions obtained from the embedded LP problem (7). These are returned by Algorithm 2, which uses an optimal basis of the LP problem to calculate them. Inspiration for Algorithm 2 came from a related algorithm for solving ODEs containing embedded LP problems [13], e.g. DFBA models. However, significant differences between DFBA (continuous ODEs) and SSA (discrete events) necessitates a distinct, novel methodology and software implementation. It should be noted that our algorithm is compatible with lexicographic optimisation for LP uniqueness as also described in [13] in order to address the problem that solutions to the FBA problem (propensity values of SSA-FBA reactions) are not necessarily unique. Algorithm 1 is outlined below, and Algorithm 2 will be described after presenting the necessary overview of optimal bases in LP problems.

Algorithm 1 depends on the reaction probability density function

$$p(\tau, j|\mathbf{n}^s, t) = \bar{a}_j(\mathbf{n}^s)\exp\left(-\bar{a}_0(\mathbf{n}^s)\tau\right), \tag{8}$$

where

$$\bar{a}_0(\mathbf{n}^s) = \sum_{j \in \mathscr{R}} \bar{a}_j(\mathbf{n}^s). \tag{9}$$

In Algorithm 1, propensity values for SSA only reactions are calculated by direct evaluation of the propensity function, whereas propensity values for SSA-FBA reactions are calculated by solving the embedded LP problem in Algorithm 2. Propensity values for FBA only reactions are not required for simulation of an SSA-FBA model because they do not appear in the MCME (2). Therefore, the subset $\mathscr{R}$ in (9) contains the SSA only and SSA-FBA reaction indices, excluding those of FBA only reactions. An informed reader will notice the similarity between Algorithm 1 and the slow-scale SSA algorithm [7, 14]. Indeed, Algorithm 1 is nearly identical with the main difference being the use of Algorithm 2 to calculate propensity values for SSA-FBA reactions. In slow-scale SSA however, where the goal is to provide a good approximation of trajectories of the MCME (2) rather than accommodate large metabolic network models lacking kinetic information, expected propensities are approximated using moments of an approximation of $P(\mathbf{n}^f|\mathbf{n}^s)$ [3, 7].

Before describing Algorithm 2, we briefly recall the use of (non-)basic variables for solving LP problems such as (7) that take the more general form

$$\begin{aligned} &\text{maximise}: \quad z = \mathbf{c} \cdot \mathbf{a}_S \\ &\text{subject to}: \quad \mathbf{a}_R = \mathbf{A} \cdot \mathbf{a}_S, \quad \mathbf{l}_R(t) \le \mathbf{a}_R \le \mathbf{u}_R(t), \quad \mathbf{l}_S(t) \le \mathbf{a}_S \le \mathbf{u}_S(t). \end{aligned} \tag{10}$$

Here maximisation is performed with respect to the *structural* variables $\mathbf{a}_S$ corresponding to propensity values $\bar{\mathbf{a}}_{FBA}$ in the original LP problem (7). The problem can be written compactly as

$$\begin{aligned} &\text{maximise}: \quad z = (\mathbf{0}, \mathbf{c}) \cdot \mathbf{a} \\ &\text{subject to}: \quad (\mathbf{I}, -\mathbf{A}) \cdot \mathbf{a} = 0, \quad \mathbf{L}(t) \le \mathbf{a} \le \mathbf{U}(t), \end{aligned} \tag{11}$$

**Algorithm 1** Simulation of an SSA-FBA model

Initialise: Set $t = 0$, initial state $\mathbf{n}^s(0)$, $j = 0$, and fix a simulation end time $T$

1. Given $j$ and the current state $\mathbf{n}^s(t)$: compute SSA only propensity values by evaluating rate equations, use Algorithm 2 to obtain propensity values for SSA-FBA reactions, and calculate $\bar{a}_0$ in (9).

2. Calculate the stochastic time step as

$$\tau = -\frac{1}{\bar{a}_0} \ln r_1$$

   where $r_1$ is a uniformly distributed number on $[0,1]$

3. Sample a second random number $r_2$ uniformly distributed on $[0,1]$ and set $j$ such that

$$\sum_{i \in \mathscr{R}_{<j}} \frac{\bar{a}_i}{\bar{a}_0} \leq r_2 \leq \sum_{i \in \mathscr{R}_{\leq j}} \frac{\bar{a}_i}{\bar{a}_0}$$

   where $\mathscr{R}_{<j}$, $\mathscr{R}_{\leq j}$ are the subsets of $\mathscr{R}$ containing all reaction indices $i : j > i \in \mathscr{R}$ and $i : j \geq i \in \mathscr{R}$, respectively.

4. Update the number of slow species according to $\mathbf{n}^s(t+\tau) = \mathbf{n}^s(t) + \mathbf{S}^s_j$ and let $t \leftarrow t + \tau$. If $t \geq T$ stop simulation, otherwise return to Step 1.

**Algorithm 2** Return current SSA-FBA propensity values

Parameters: $j \in \mathscr{R}$, $\mathbf{n}^s(t)$ from Algorithm 1, *basis_valid* = `True`, and *dependency_list* $= \varnothing$.

1. If $j = 0$, update all bounds using $\mathbf{n}^s(0)$ and solve the LP problem (7) using the Simplex Method. Calculate resulting propensity values for all SSA-FBA reactions and store the simplex tableau matrix, current bound values, and the current values of basic variables. Otherwise, if $j > 0$, find all $\bar{a}_{FBA,i}$ corresponding to SSA-FBA and FBA only reactions with bounds that depend on slow species produced/consumed by reaction $j$ and do the following:

   1.1. For each such basic variable $\bar{a}_{FBA,i}$, update its bound with the new supplied value and add its index $i$ to the set *dependency_list*

   1.2. For each such non-basic variable $\bar{a}_{FBA,i}$ not fixed at the relevant bound, update its bound with the new supplied value

   1.3. For each such non-basic variable $\bar{a}_{FBA,i}$ fixed at the relevant bound, update its bound with the new supplied value and identify all basic variables corresponding to non-zero entries in the respective column of the simplex tableau matrix. For each $k$th such basic variable, add to its current value the difference between the new and previous bound value (weighted by the corresponding entry of the simplex tableau matrix) and enter its index $k$ to the set *dependency_list*

2. If $j = 0$, skip this step. Otherwise, if $j > 0$, check that all basic variables in *dependency_list* remain within their new bounds. If true, keep the propensity values of the SSA-FBA reactions using either stored basic values (if the propensity value is basic) or the current bound values (if the propensity value is non-basic). If false, re-solve the LP problem (7) using the Simplex Method, calculate the resulting propensity values for all SSA-FBA reactions, and store the simplex tableau matrix and the current values of the basic variables. Calculate the propensity values for SSA-FBA reactions using either the stored basic values (if the propensity value is basic) or the current bound values (if the propensity value is non-basic).

3. Return previously calculated propensity values for all SSA-FBA reactions

where $\mathbf{a} = (\mathbf{a}_R, \mathbf{a}_S)^T$ is the augmented set of variables also containing the *auxiliary* variables $\mathbf{a}_R$. The constraint matrix $\mathbf{A}$ has dimension $d_R \times d_S$, where $d_R$ is the number of auxiliary variables, $d_S$ the number of structural variables, and bound vectors $\mathbf{L}(t) = (\mathbf{l}_S(t), \mathbf{l}_R(t))^T$ and $\mathbf{U}(t) = (\mathbf{u}_S(t), \mathbf{u}_R(t))^T$ both have dimension $d_S + d_R$. We refer to the LP problem (11) at time $t$ as $LP(t)$, where the only difference between $LP(t_1)$ and $LP(t_2)$ are different values of the upper and lower bounds at time points $t_1$ and $t_2$. This representation is also useful because by using it one does not distinguish between structural and auxiliary variables.

A variable is called *non-basic*, if its (lower or upper) bound is active, i.e. the variable is fixed at that bound; otherwise it is called *basic*. In a *basic solution*, there are always $d_S$ non-basic variables and $d_R$ basic variables, which corresponds to the situation in which exactly $d_R$ bounds are active. It is a well-known result in the theory of LP problems that if (11) has an optimal solution, it has (at least one) *basic feasible solution* that maximises the objective $z$. Here *feasible* implies that the basic solution lies within the constraints of the LP problem (11). In a basic solution, the $d_R$-dimensional vector $\mathbf{a}_B(t)$ of basic variable values is related to the $d_S$-dimensional vector $\mathbf{a}_N(t)$ of non-basic variables fixed at either their upper or lower bound (hence explicit $t$-dependence) by the $d_R \times d_S$ simplex tableau matrix $\Xi$:

$$\mathbf{a}_B(t) = \Xi \cdot \mathbf{a}_N(t) \tag{12}$$

where the augmented vector $(\mathbf{a}_B, \mathbf{a}_N)^T = \Pi \cdot \mathbf{a}$ is related to $\mathbf{a}$ by a permutation matrix $\Pi$.

In the fast SSA-FBA simulation algorithm, the Simplex Method is first used to obtain an optimal basic feasible solution of the form (12) for the initial problem $LP(0)$ at the beginning of a simulation. The Simplex Method defines an associated simplex tableau matrix $\Xi$ and the set of active constraints, i.e., which non-basic variables $\mathbf{a}_N(0)$ are fixed at which initial bounds $\mathbf{L}(0), \mathbf{U}(0)$. After a reaction event is executed at time $t + \tau$, determining an optimal basic feasible solution to the new problem $LP(t + \tau)$ requires either invoking the Simplex Method or using Equation (12) with the simplex tableau matrix $\Xi$ from a solution to the previous problem $LP(t)$. The decision for how the new optimal basic feasible solution is calculated by Algorithm 2 is based on the following claim.

**Claim 1.** Given an optimal basic feasible solution $(\mathbf{a}_B(t), \mathbf{a}_N(t))$ of LP problem $LP(t)$ with associated simplex tableau matrix $\Xi$, the solution

$$\mathbf{a}_B(t+\tau) = \Xi \cdot \mathbf{a}_N(t+\tau), \tag{13}$$

where the arrangement of active constraints of non-basic variables $\mathbf{a}_N(t+\tau)$ remains unchanged, is an optimal basic feasible solution to problem $LP(t+\tau)$ if basic variables $\mathbf{a}_B(t+\tau)$ remain within their new bounds.

*Proof.* Proving this claim depends on showing that the Karush-Kahn-Tucker (KKT) conditions for an optimal basic feasible solution to $LP(t+\tau)$ are satisfied as long as basic variables $\mathbf{a}_B(t+\tau)$ remain within the new set of bounds $\mathbf{L}(t+\tau), \mathbf{U}(t+\tau)$. The KKT conditions are

1. $(\mathbf{I}, -\mathbf{A}) \cdot \mathbf{a} = 0$
2. $(\mathbf{I}, -\mathbf{A})^T \cdot \boldsymbol{\pi} + \boldsymbol{\lambda}_L + \boldsymbol{\lambda}_U = (\mathbf{0}, \mathbf{c})^T$
3. $\mathbf{L}(t+\tau) \leq \mathbf{a} \leq \mathbf{U}(t+\tau)$
4. $\boldsymbol{\lambda}_L \leq \mathbf{0}$, $\boldsymbol{\lambda}_U \geq \mathbf{0}$
5. $\mathbf{D}(\boldsymbol{\lambda}_L) \cdot (\mathbf{a} - \mathbf{L}(t+\tau)) = \mathbf{D}(\boldsymbol{\lambda}_U) \cdot (\mathbf{a} - \mathbf{U}(t+\tau)) = \mathbf{0}$

where $\boldsymbol{\pi}$ and $\boldsymbol{\lambda}_U, \boldsymbol{\lambda}_L$ are a $d_R$- and $(d_S + d_R)$-dimensional vectors, respectively, of Lagrange multipliers. $\mathbf{D}$ is the operator that takes a $d$-dimensional vector to a square $(d \times d)$-dimensional matrix whose only nonzero entries are the vector's components ordered along the diagonal. The claim assumes that $LP(t)$ has an optimal basic feasible solution with associated simplex tableau matrix $\Xi$ and therefore (13) satisfies Condition 1 by construction because these constraints do not change in $LP(t+\tau)$. Similarly, because Condition 2 is identical for both problems $LP(t)$ and $LP(t+\tau)$, and the set of active constraints for non-basic variables $\mathbf{a}_N(t+\tau)$ remains unchanged (perhaps with new upper or lower bound values), Lagrange multipliers $\boldsymbol{\pi}$ and $\boldsymbol{\lambda}_U, \boldsymbol{\lambda}_L$ are free to take on the same values as those in problem $LP(t)$ in order to satisfy Conditions 2, 4, 5. This implies that only Condition 3 must be validated for (13) to be an optimal basic feasible solution to problem $LP(t+\tau)$, and since non-basic variables must be fixed at their new bounds this remaining condition is guaranteed to hold if basic variables $\mathbf{a}_B(t+\tau)$ do not exceed them either. □

Claim 1 justifies the method used for numerical calculation of SSA-FBA propensity values in Algorithm 2. After a reaction event has been executed in Algorithm 1 and the resulting counts of slow species used to update problem $LP(t+\tau)$, Equation (13) generates basic variables that are evaluated for violation of their bounds. Whenever a basic variable is found to lie outside its bounds, the Simplex Method is called to obtain a new optimal basic feasible solution and an associated simplex tableau matrix for $LP(t+\tau)$. However, if all basic variables remain within their bounds then (13) provides the full set of numerical values from which propensity values for SSA-FBA reactions can be returned and, consequently, the number of times the Simplex Method is called over the course of a simulation can be reduced dramatically.

In addition to evaluating the approximate SSA-FBA simulation method using the toy model described in the main text (see Figures 3a-c in main text for results), we evaluated the performance of our fast SSA-FBA simulation method using the *Mycoplasma genitalium* metabolic network model by simulating random sequences of (50000, 100000, or 200000) reaction execution events. Execution of each individual event corresponded to a random selection of one to five SSA-FBA or FBA only reactions whose upper bound values are known in the original *M. genitalium* metabolic model (i.e., not zero or infinity), and reducing their upper bounds through multiplication by a constant less than one (0.9 used in experiments presented here). The cumulative effect was to slowly drive the growth rate of *M. genitalium* to zero over the course of simulation. After execution of each event, SSA-FBA and FBA only propensity values were calculated either by directly solving the embedded FBA problem or using the fast optimal basis algorithm. Employing this approach rather than evaluating a full SSA-FBA simulation allowed us to confidently compare the performance of direct and fast methods, while confirming that calculated propensity values remained identical. Figure 3d in the main text displays representative results of the comparisons between the fast and direct SSA-FBA simulation methods, showing that the former improves run time by an order of magnitude over the latter. The fast optimal basis algorithm also scales better in the number of reaction execution events than the direct implementation (slopes of linear regressions using data from Figure 3d are 19.0 versus 1.7 for direct versus fast, respectively).

An SSA-FBA simulation package implementing the fast, direct and approximate SSA-FBA simulation methods for generic SSA-FBA models is available for free at https://gitlab.com/davidtourigny/single-cell-fba. The package is distributed as a Python extension module written in C++ and includes code that enables users to reproduce the tests, toy model and single-cell *M. pneumoniae* model described here and in the main text.

### S3. Extended description of reduced single-cell model

To enable us to use a model to gain insights into the molecular underpinnings of the metabolism of individual *Mycoplasma pneumoniae* cells with modest complexity, we decided to construct a model that could explain the growth of *M. pneumoniae* cells and account for the functional contribution of each individual gene to growth. Consequently, we chose to construct a model that represents the nutrient import and export, metabolism, transcription, translation, macromolecular complexation, and RNA and protein turnover of *M. pneumoniae* and the genes which catalyse these functions. We simulated the model using SSA-FBA, with metabolism represented using FBA and the non-metabolic processes represented using SSA.

Here, we outline some key features of the model. The full details of the model are defined in Supplementary File S1. Supplementary File S1 is also the input file to the script for simulating models found within the SSA-FBA simulation package available for free at https://gitlab.com/davidtourigny/single-cell-fba. The script uses this file to represent the model as an SSA-FBA model and uses SSA-FBA to simulate it. Alternative models can be built by modifying this file. Supplementary File S1 defines each compartment, species, reaction, rate law, rate parameter, and initial condition of the model.

#### Compartments represented by the model

To capture *M. pneumoniae* cells and their external environment as simply as possible, we decided that the model should represent two compartments: *M. pneumoniae* cells, including their cytosols and membranes, and the external environment outside their membranes.

#### Species and reactions represented by the model

The exact scope of the model was driven by the genomic sequence of *M. pneumoniae* [16], its annotation [23, 24], its reconstructed metabolism [15], its genetic code (https://www.ncbi.nlm.nih.gov/Taxonomy/Utils/wprintgc.cgi), and its reconstructed complexome [25, 26, 19]. First, we determined the biochemical processes that the model must represent, including the transcription of each RNA; the translation of each protein; the assembly of each complex; the turnover of each RNA, protein, and complex; and the metabolic reactions needed to synthesise the substrates of these processes and recycle the byproducts of these processes, namely the reactions needed to produce nucleotide triphosphates, amino acids, and water for transcription and translation, and the reactions needed to recycle nucleotide diphosphates, nucleotide monophosphates, diphosphates, phosphates, hydrogen ions, and amino acids produced by transcription, translation, and the turnover of RNAs, proteins, and complexes.

In turn, this established the enzymes that the model must represent, namely the genes required to transcribe RNAs (RNA polymerase), translate proteins (ribosome), turnover RNAs (RNases), turnover proteins (proteases), turnover complexes (RNases and proteases), and produce and recycle the metabolites described above (metabolic enzymes). The metabolic enzymes required for these metabolic functions were determined using the reconstructed metabolism of *M. pneumoniae* [15]; specifically, we used the reconstruction to determine the minimum set of enzymes required for these metabolic functions.

Subsequently, this established the genes that the model must represent, namely the genes that code for each enzyme. We used the annotation of the genome of *M. pneumoniae*, the gene-reaction rules of the metabolic reconstruction of *M. pneumoniae* [15], and the reconstructed complexome of *M. pneumoniae* [25, 26, 19] to determine the genes that code for each enzyme.

Together, this established the species that the model must represent, including a species for each RNA transcript of each gene, a species for each protein of each protein-coding gene, a species for each macromolecular complex, and species for each metabolite involved in each metabolic reaction. We determined the sequence of each RNA species using the genomic sequence of *M. pneumoniae* and the start and stop coordinate of each gene. We determined the sequence of each protein species using the sequence of its RNA template and the genetic code of *M. pneumoniae* (https://www.ncbi.nlm.nih.gov/Taxonomy/Utils/wprintgc.cgi). We obtained the structure of each complex from previous reconstructions [25, 26, 19]. We obtained the structure of each metabolite from the previous metabolic reconstruction [15]. We used BpForms and BcForms [30] to calculate the chemical formula and molecular weight of each RNA, protein, and complex species.

The model represents 93 genes (13% of the genome of *M. pneumoniae*), 293 species, and 506 reactions. This includes 83 metabolites, 93 RNAs, 90 proteins, 27 complexes, 33 enzymes, 86 metabolic reactions, 93 transcription reactions, 90 translation reactions, 27 complex assembly reactions, 93 RNA turnover reactions, 90 protein turnover reactions, and 27 complex turnover reactions. In addition, the metabolism portion of the model involves one pseudo-species that represents the biomass of the cell, one pseudo-reaction that represents its synthesis, and 32 exchange pseudo-reactions for each extracellularly-localized metabolite.

#### Equations and rate laws of the non-metabolic reactions

The rate laws of the non-metabolic reactions were chosen to capture the effect of the concentration of each substrate and enzyme on the rate of each reaction. In particular, each rate law was chosen to be the product of the maximum turnover rate of each enzyme, logistic terms of the concentration of each substrate, and a linear term of the concentration of the enzyme. The logistic terms capture the behaviours that the rates of reactions vanish when no substrate is present and that the rates of reactions saturate at high substrate concentrations when each enzyme is occupied.

##### Transcription

The model represents the transcription of each RNA $i$ as a single lumped reaction that fuses nucleotide triphosphates, $\mathrm{NTP}_j$ into a sequence-dependent polymer, where $m_{ij}$ represents the count of nucleotide $j$ in RNA $i$

$$\sum_j m_{ij}\mathrm{NTP}_j + \mathrm{H}^+ \rightarrow \mathrm{RNA}_i + \left(\sum_j m_{ij}\right)\mathrm{PPi}. \tag{14}$$

The rate, $v_{\mathrm{r},i}^{\mathrm{syn}}$ of the transcription of each RNA $i$ is modelled as the product of the maximum rate of its transcription $k_{\mathrm{r},i}^{\mathrm{syn}}$, logistic terms of the concentration of each nucleotide triphosphate, $M_j$, and the concentration of RNA polymerase, $E_{\mathrm{pol}}$

$$v_{\mathrm{r},i}^{\mathrm{syn}} = k_{\mathrm{r},i}^{\mathrm{syn}}\left(\prod_j \frac{M_j}{K_{\mathrm{M,n}} + M_j}\right)E_{\mathrm{pol}}. \tag{15}$$

##### Translation

Similarly, the model represents the translation of each protein $i$ as a single lumped reaction powered by the hydrolysis of GTP that fuses amino acids, $\mathrm{AA}_j$ into a sequence-dependent polymer, where $m_{ij}$ represents the count of amino acid $j$ in protein $i$

$$\begin{gathered}\sum_j m_{ij}\mathrm{AA}_j + \left(\sum_j 2m_{ij} + 3\right)\mathrm{GTP} + \left(\sum_j m_{ij} + 4\right)\mathrm{H_2O} \rightarrow \\ \mathrm{protein}_i + \left(\sum_j 2m_{ij} + 3\right)\left(\mathrm{GDP} + \mathrm{Pi} + \mathrm{H}^+\right).\end{gathered} \tag{16}$$

The rate, $v_{\mathrm{p},i}^{\mathrm{syn}}$ of the translation of each protein $i$ is modelled as the product of the maximum rate of its translation $k_{\mathrm{p},i}^{\mathrm{syn}}$, logistic terms of the concentration of each amino acid, $M_j$, and the concentration of its template RNA, $R_i$, and the concentration of ribosomes, $E_{\mathrm{ribo}}$

$$v_{\mathrm{p},i}^{\mathrm{syn}} = k_{\mathrm{p},i}^{\mathrm{syn}}\left(\prod_j \frac{M_j}{K_{\mathrm{M,a}} + M_j}\right)\frac{R_i}{K_{\mathrm{M,r}_i} + R_i}E_{\mathrm{ribo}}. \tag{17}$$

##### Macromolecular complexation

The model represents the synthesis of each complex $i$ as a single lumped reaction that agglomerates $r_{ij}$ molecules of each RNA subunit $j$ and $p_{ij}$ molecules of each protein subunit $j$

$$\sum_j r_{ij}\text{RNA}_j + \sum_j p_{ij}\text{protein}_j \rightarrow \text{complex}_i. \tag{18}$$

The rate, $v^{\text{syn}}_{\text{c},i}$ of the assembly of each complex $i$ is modelled as the product of the maximum rate of its assembly $k^{\text{syn}}_{\text{c},i}$ and the minimum concentration of its subunits, $R_j$ and $P_j$

$$v^{\text{syn}}_{\text{c},i} = k^{\text{syn}}_{\text{c},i} \min\left(\min_{j \text{ st. } r_{ij}>0} R_j, \min_{j \text{ st. } p_{ij}>0} P_j\right). \tag{19}$$

##### RNA degradation

The model represents the hydrolytic degradation of each RNA $i$ as a single lumped reaction that disassembles polymers to individual nucleotide monophosphates, $\text{NMP}_j$, where $m_{ij}$ represents the count of nucleotide $j$ in RNA $i$

$$\text{RNA}_i + \left(\sum_j m_{ij} - 1\right)\text{H}_2\text{O} \rightarrow \sum_j m_{ij}\text{NMP}_j + \left(\sum_j m_{ij} - 1\right)\text{H}^+. \tag{20}$$

The rate, $v^{\text{deg}}\text{r}, i$ of the degradation of each RNA $i$ is modelled as the product of the maximum rate of its degradation $k^{\text{deg}}_{\text{r},i}$, a logistic term of the concentration of the RNA, $R_i$, and the concentration of oligoribonuclease NrnA, $E_{\text{NrnA}}$

$$v^{\text{deg}}_{\text{r},i} = k^{\text{deg}}_{\text{r},i} \frac{R_i}{K_{\text{M},\text{r}_i} + R_i} E_{\text{NrnA}}. \tag{21}$$

##### Protein degradation

Similarly, the model represents the hydrolytic degradation of each protein $i$ as a single lumped reaction that disassembles polymers to individual amino acids, $\text{AA}_j$, where $m_{ij}$ represents the count of amino acid $j$ in protein $i$

$$\text{protein}_i + \left(\sum_j m_{ij} - 1\right) \text{H}_2\text{O} \rightarrow \sum_j m_{ij} AA_j. \tag{22}$$

The rate, $v_{\text{p},i}^{\text{deg}}$ of the degradation of each protein $i$ is modelled as the product of the maximum rate of its degradation $k_{\text{p},i}^{\text{deg}}$, a logistic term of the concentration of the protein, $P_i$, and the concentration of protease Lon, $E_{\text{Lon}}$

$$v_{\text{p},i}^{\text{deg}} = k_{\text{p},i}^{\text{deg}} \frac{P_i}{K_{\text{M,p}_i} + P_i} E_{\text{Lon}}. \tag{23}$$

**Complex degradation**

Similarly, the model represents the hydrolytic degradation of each complex $i$ as a single lumped reaction that disassembles polymers to individual nucleotide monophosphates, $\text{NMP}_j$, and amino acids, $\text{AA}_j$, where $m_{\text{n},ij}$ and $m_{\text{a},ij}$ represent the counts of nucleotide monophosphates and amino acids $j$ in complex $i$

$$\begin{gathered}\text{complex}_i + \left(\sum_j m_{\text{n},ij} + \sum_j m_{\text{a},ij} - 2\right) \text{H}_2\text{O} \rightarrow \\ \sum_j m_{\text{n},ij} \text{NMP}_j + \sum_j m_{\text{a},ij} AA_j + \left(\sum_j m_{\text{n},ij} - 1\right) \text{H}^+.\end{gathered} \tag{24}$$

The rate, $v_{\text{c},i}^{\text{deg}}$ of the degradation of each complex $i$ is modelled as the product of the maximum rate of its degradation $k_{\text{c},i}^{\text{deg}}$, a logistic term of the concentration of the complex, $C_i$, and the concentration of protease Lon, $E_{\text{Lon}}$

$$v_{\text{c},i}^{\text{deg}} = k_{\text{c},i}^{\text{deg}} \frac{C_i}{K_{\text{M,c}_i} + C_i} E_{\text{Lon}} \tag{25}$$

#### Flux bounds of the model

To capture the impact of gene expression on metabolism, we chose to bound the flux of metabolic reactions by the product of the concentration of its enzyme and the maximum turnover rate ($k_{cat}$) of the enzyme. Unfortunately, experimental measurements of these values are not available for *M. pneumoniae*, but many are for other bacterial species in the BRENDA database [27]. If the measured $k_{cat}$ of a given enzyme was available for one or more wild-type bacterial species in BRENDA. We used the largest of these values (multiplied by the concentration of enzyme) to bound the FBA reaction catalysed by that enzyme as described above. Conversely, FBA reactions catalysed by enzymes for which no measured $k_{cat}$ values are available in BRENDA were left unbounded. An exception to this rule was that the same $k_{cat}$ values were used for reactions differing only by the use of a cofactor (e.g., $k_{cat}$ value available for reaction catalysed by enzyme using ATP was also used as $k_{cat}$ value for the same enzyme using GTP if kinetic data for the latter are not available).

#### Biosynthetic pseudo-reaction of the model

Typically, flux-balance analysis models include a pseudo-reaction that represents the average molecular composition of cells and the average energy required to assemble and maintain cells. Similar to our previous work [19], our model generalizes this reaction to encompass the substrates that the metabolic machinery must produce to support all of the other processes in a cell, as well as the byproducts of these processes that the metabolic machinery must recycle. For this model, the biomass synthesis pseudo-reaction represents the production of nucleotide triphosphates, amino acids, and water for transcription, translation and RNA and protein turnover and the recycling of nucleotide diphosphates, nucleotide monophosphates, amino acids, phosphates, diphosphates, and hydrogen ions produced by transcription; translation; and the turnover of RNAs, proteins, and complexes.

#### Parameters of the rate laws and initial conditions

The model includes 1,003 parameters. This includes 33 parameters for the mean initial intracellular count of each intracellular metabolite, 32 parameters for the mean initial extracellular count of each extracellular metabolite, 93 parameters for the mean initial intracellular count of each RNA, 90 parameters for the mean initial intracellular count of each protein, 27 parameters for the mean initial intracellular count of each complex, 27 parameters for the maximum turnover rates of 27 metabolic reactions (described in previous section "Flux bounds of the model"), 93 parameters for the maximum synthesis rate of each RNA, 90 parameters for the maximum synthesis rate of each protein, 1 parameter for the maximum assembly rate of the complexes, 93 parameters for the maximum turnover rate of each RNA, 90 parameters for the maximum turnover rate of each protein, 27 parameters for the maximum turnover rate of each complex, 1 parameter for the affinity of RNA polymerase for nucleotide triphosphates, 90 parameters for the affinity of ribosomes for each mRNA, 1 parameter for the affinity of ribosomes for amino acids, 93 parameters for the affinity of RNases for each RNA, 90 parameters for the affinity of proteases for each protein, 27 parameters for the affinity of proteases for each complex, 2 parameters for the fluxes of the import and export of nutrients, 1 parameter for the average initial volume of each cell, and 1 parameter for the initial density of cells in their growth media.

We used several types of experimental biochemical and physiological data to calibrate the model to represent the growth of *M. pneumoniae*. This includes data about the sequence of the *M. pneumoniae* genome [16]; the start and stop coordinates and direction of each gene in *M. pneumoniae* [23, 24]; the typical concentrations of metabolites in bacteria [20]; the abundances of *M. pneumoniae* RNAs [29] and proteins [28]; the half-lives of *M. pneumoniae* RNAs and proteins [28]; the turnover rates of 27 metabolic reactions in various bacteria [27], the fluxes of the import and export of nutrients by the closely related bacterium *Mycoplasma genitalium* [21]; the average volume of *M. pneumoniae* cells [17]; the typical water content of bacteria [22]; the chemical composition of the Hayflick medium typically used to culture *M. pneumoniae* (PPLO media, Becton, Dickinson and Company; Beef heart infusion A1502, Solabia Group; Donor horse serum, Corning; Pork meat peptone - A1728, Solabia Group) and that of SP4 media frequently used to culture *M. genitalium* [19]; and the density of *M. pneumoniae* cells in cultures [15].

##### Distributions of the initial count of each species

First, we recognized that the mean initial abundance of each species within individual cells is approximately equal to the population-average abundance of each species, which has been experimentally characterized. Next, we approximated the mean initial abundance of each complex as the minimum ratio of the total experimentally observed abundance of each subunit [29, 28] to its reconstructed stoichiometry within the complex [25, 26, 19]. Third, we approximated the mean initial free abundance of each RNA and protein as the difference between its total observed abundance and the mean initial abundance estimated to be bound to complexes. For proteins that do not participate in complexes, the free abundance is equal to its total observed abundance.

We estimated the mean initial intracellular concentration of each metabolite as its observed population-average concentration, equal to 1 mM for each nucleotide triphosphate, diphosphate, and monophosphate; 0.5 mM for each amino acid; 1 mM for diphosphate; 5 mM for phosphate 11.2 nM (equal to pH 7.75) for hydrogen ions; and 55 M for water. We estimated the mean initial extracellular concentration of each metabolite similarly. To represent the physiology of individual cells, the initial conditions of the model describe the distribution of the count of each species in cells and in their growth medium. The distribution of the count of each species is modelled as a Poisson distribution with a mean equal to the average value of that species across the cycle. At the beginning of each simulation, these distributions are sampled to determine the initial conditions of the simulation.

##### Rate parameters of the non-metabolic reactions

Following the rule of thumb that the affinity of an enzyme for a substrate is often similar to the concentration of the substrate, first, we approximated each $K_{\mathrm{M}}$ as the average experimentally observed concentration of its associated substrate. Specifically, we equated $K_{\mathrm{M,n}}$ of Equation 15 to

1 mM [20]; we equated $K_{\mathrm{M,a}}$ of Equation 17 to 0.5 mM [20]; and we equated $K_{\mathrm{M,r}_i}$, $K_{\mathrm{M,p}_i}$, and $K_{\mathrm{M,c}_i}$ of Equations 17, 21, 23, and 25 to the average abundance of each RNA, protein, and complex estimated in the previous subsection.

Next, we equated the modelled degradation rate of each RNA, $v^{\mathrm{deg}}_{\mathrm{r},i}$, to the observed degradation rate equal to $ln(2)/\tau_i R_i$, where $\tau_i$ is the experimentally observed half-life of RNA $i$. Using the value of $K_{\mathrm{M,r}_i}$ estimated above and rearranging yields $k^{\mathrm{deg}}_{\mathrm{r},i} = 2ln(2)/\tau_i R_i/E_{\mathrm{NrnA}}$. A similar analysis for the degradation of proteins and complexes yields $k^{\mathrm{deg}}_{\mathrm{p},i} = 2ln(2)/\tau_i P_i/E_{\mathrm{Lon}}$ and $k^{\mathrm{deg}}_{\mathrm{c},i} = 2ln(2)/\tau_i C_i/E_{\mathrm{Lon}}$.

Finally, to capture the growth of *M. pneumoniae* cells, the average rate of synthesis of each RNA, protein, and complex must be balanced by the average rate of its degradation. This implies that $v^{\mathrm{syn}}_{x,i} = v^{\mathrm{deg}}_{x,i}$ for $x \in \{\mathrm{r},\mathrm{p},\mathrm{c}\}$. Using the values of $v^{\mathrm{deg}}_{x,i}$ and $K_{\mathrm{M,y}_i}$ estimated above yields $k^{\mathrm{syn}}_{r,i} = 2^4 ln(2)/\tau_i R_i/E_{\mathrm{pol}}$ and $k^{\mathrm{syn}}_{p,i} = 2^{21} ln(2)/\tau_i P_i/E_{\mathrm{ribo}}$. We approximated the assembly of each complex as a fast process with $k^{\mathrm{syn}}_{c,i}$ as $1/60\,\mathrm{s}^{-1}$.

**Coefficients of the biosynthetic pseudo-reaction**

To capture the growth of *M. pneumoniae* cells, the average output of the modelled metabolism of *M. pneumoniae* must be the inverse of the average output of the other modelled processes. Specifically, the average rate of metabolic production of nucleotide triphosphates must equal the average rate of consumption of nucleotide triphosphates by the other modelled processes. Similarly, the average rate of metabolic recycling of nucleotide diphosphates, nucleotide monophosphates, diphosphate, and phosphate must equal the average rate of production of these metabolites by the other modelled processes.

Therefore, we estimated the coefficient of each metabolite of the biosynthetic pseudo-reaction as its total average rate of consumption or production across all of the other modelled processes. We estimated the average rate of consumption/production of each metabolite in each non-metabolic process by (a) evaluating the rate laws (Equations 15, 17, 17, 19, 23, and 25) with the average abundance of each species estimated above and (b) multiplying each result by the stoichiometry of the metabolite in the corresponding reaction.

#### Annotation of the semantic meaning of each species and reaction

These precise semantic meaning of each species and reaction is defined in Supplementary File S1. The semantic meaning of each metabolite species is defined using SMILES and annotated using a ChEBI or PubChem identifier. The semantic meaning of each RNA species is defined by its genomic coordinates and direction and annotated using its sequence and an NCBI Gene identifier. The semantic meaning of each protein species is defined by the sequence of its RNA transcript and annotated using its sequence and a UniProt identifier. The semantic meaning of each complex is defined by the stoichiometries of its subunits. Supplementary File S1 also describes the chemical formula, molecular weight, and charge of each species. The semantic meaning of each reaction is defined by the stoichiometry of each substrate and product. Where applicable, the semantic meaning of the metabolic reactions is also annotated using Enzyme Commission (EC) numbers.

#### Additional details of simulation

Simulations of the model used the SSA-FBA simulation package available at https://gitlab.com/davidtourigny/single-cell-fba. As described at the end of Supplementary Appendix S1, the relative scale of SSA only to SSA-FBA propensity values (based on the stoichiometry of SSA-FBA reactions) serves as an additional parameter that controls the degree to which the simulation approximates biology and and can dictate the speed of simulation. For simulations described in the main text, the relative stoichiometry of SSA-FBA reactions was chosen to be 200, implying that execution of an SSA-FBA reaction event corresponds to updating species count by 200 times the stoichiometry of the corresponding SSA-FBA selected reaction. This in turn corresponds to a relative scaling factor of 0.005.

It is common for the population of some species to be driven negative values in hybrid deterministic and stochastic adaptations of SSA [31]. In order to prevent unphysiological negative species counts from arising in simulations, the SSA-FBA simulation package https://

gitlab.com/davidtourigny/single-cell-fba employs the Zero-Reaction rule described in [31] as the best remedy for nonlinear and sensitive systems considering its efficiency and simplicity.

To remove internal cycles from optimal solutions to the embedded FBA problem (7) we employed lexicographic optimisation using a second, parsimonious FBA objective (minimising the sum of absolute flux values). Briefly, after solving (7), the resulting optimal value $z^*$ is used to bound the objective function as an additional constraint in the LP problem

$$\begin{aligned} &\text{minimise}: \quad ||\bar{\mathbf{a}}_{FBA}||_1 \\ &\text{subject to}: \quad \mathbf{S}^f \cdot \bar{\mathbf{a}}_{FBA} = 0, \quad \mathbf{l}(t) \leq \bar{\mathbf{a}}_{FBA} \leq \mathbf{u}(t), \quad \mathbf{c} \cdot \bar{\mathbf{a}}_{FBA} \geq z^* \end{aligned} \tag{26}$$

where $||\mathbf{y}||_1 = \sum_{k=1}^{N} |y_k|$ denotes the $L_1$ norm of a vector $\mathbf{y} = (y_1, y_2, ..., y_N)$. Subsequently, the optimal solution to (26) is used to return propensity values for the SSA-FBA reactions in the model at time $t$. Although lexicographic optimisation is in principle compatible with the optimal basis algorithm, the implementation of parsimonious FBA in https://gitlab.com/davidtourigny/single-cell-fba is provided for the direct method only.
